# A Chemical Reaction Similarity-Based Prediction Algorithm Identifies the Multiple Taxa Required to Catalyze an Entire Metabolic Pathway of Dietary Flavonoids

**DOI:** 10.1101/2023.05.05.539480

**Authors:** Ebru Ece Gulsan, Farrhin Nowshad, Pomaikaimaikalani Yamaguchi, Xiaokun Dong, Arul Jayaraman, Kyongbum Lee

**Affiliations:** Tufts University; Texas AM University

## Abstract

Flavonoids are polyphenolic phytochemicals abundant in plant-based, health-promoting foods. They are only partially absorbed in the small intestine, and gut microbiota plays a significant role in their metabolism. As flavonoids are not natural substrates of gut bacterial enzymes, reactions of flavonoid metabolism have been attributed to the ability of general classes of enzymes to metabolize non-natural substrates. To systematically characterize this promiscuous enzyme activity, we developed a prediction tool that is based on chemical reaction similarity. The tool takes a list of enzymes or organisms to match microbial enzymes with their non-native flavonoid substrates and orphan reactions. We successfully predicted the promiscuous activity of known flavonoid-metabolizing bacterial and plant enzymes.

Next, we used this tool to identify the multiple taxa required to catalyze an entire metabolic pathway of dietary flavonoids. Tilianin is a flavonoid-O-glycoside having biological and pharmacological activities, including neuroprotection. Using our prediction tool, we defined a novel bacterial pathway of tilianin metabolism that includes O-deglycosylation to acacetin, demethylation of acacetin to apigenin, and hydrogenation of apigenin to naringenin. We predicted and confirmed using in vitro experiments and LC-MS techniques that Bifidobacterium longum subsp. *animalis*, *Blautia coccoides* and *Flavonifractor plautii* can catalyze this pathway. Prospectively, the prediction-validation methodology developed in this work could be used to systematically characterize gut microbial metabolism of dietary flavonoids and other phytochemicals.

The bioactivities of flavonoids and their metabolic products can vary widely. We used an in vitro rat neuronal model to show that tilianin metabolites exhibit protective effect against H_2_O_2_ through reactive oxygen species (Delepine et al.) scavenging activity and thus, improve cell viability, while the parent compound, tilianin, was ineffective. These results are important to understand the gut microbiota-dependent physiological effects of dietary flavonoids.

## Introduction

Enzymes are exquisitely specific proteins that act as catalysts of biochemical reactions. They are evolved to recognize specific substrates and selectively catalyze reactions. However, some enzymes are capable of catalyzing additional reactions that may not be related to their native catalytic activities. This phenomenon is called enzyme promiscuity, which has been referred to as “underground metabolism” (Amin et al., 2019), and includes the ability of enzymes to act on substrates other than which they evolved for (Khersonsky & Tawfik, 2010). Promiscuity has a significant role in enzyme evolution and is potentially a useful tool for metabolic engineering. However, the promiscuous activity of an enzyme is not easy to detect using in vivo systems because of unknown reaction mechanisms, low levels of catalytic activities and lack of high-throughput methods to characterize enzyme-substrate pairs. Hence, promiscuous enzymes have not yet been comprehensively studied and remain to be fully utilized for metabolic engineering purposes. (Baas et al., 2013)

Flavonoids are a group of phytochemicals widely distributed in plants, fruits and vegetables that possess numerous bioactivities. They have a 15-carbon skeleton that consists of two phenyl rings (A- and B-rings) linked with a heterocyclic pyrane ring (C-ring). Depending on the substitutions on the central pyrane ring, flavonoids are classified into several subgroups: flavones, flavanones, flavonols, anthocyanins, flavan-3-ols and isoflavones. They are associated with various beneficial effects including anti-oxidant, anti-inflammatory and anti-cancer properties, making them attractive building blocks of nutraceutical, pharmaceutical and medicinal products. Their health promoting effects vary widely even among the same flavonoid subclass. (Gulsan et al., 2022)

The majority of flavonoids occur as glycosides in nature. Once flavonoids are ingested, they are partially absorbed in the small intestine, which makes them available to be metabolized by the gut microorganisms. Ingested flavonoids can be modified by removal of the sugar moiety, de/methylation and di/hydroxylation of A- and B-rings, or hydrogenation of the pyrane ring. Unlike plants, commensal gut microorganisms do not have specialized enzymes that utilize flavonoids as their native substrates. Hence, bacterial flavonoid metabolism is carried out through promiscuous enzyme activity. For example, methylation of flavonols is generally attributed to a flavonol O-methyltransferase (EC:2.1.1.76) enzyme (Schmidt et al., 2012). According to the UniProt (UniProt, 2021) and the KEGG (Kanehisa & Goto, 2000) protein databases, this enzyme can only be detected in plant species. A recent study showed that an enzyme from Streptomyces sp. KCTC 0041BP that can catalyze the methylation of quercetin, luteolin, myricetin, quercetin 3-O-β-D-glucoside and fisetin (Darsandhari et al., 2018) by cloning and heterologously expressing the enzyme in E. coli. We recently used in silico predictions and in vitro experiments to show that a chalcone synthase-like bacterial polyketide synthase can catalyze the cleavage of naringenin’s C-ring (Gülşan et al., 2022). In plants, flavanone ring cleavage is catalyzed by chalcone isomerases (Braune et al., 2016), which suggests that the polyketide synthase’s ability to cleave the C-ring of naringenin reflects a promiscuous activity. Despite the growing interest in dietary flavonoids owing to scientific evidence regarding their health-promoting effects on the host, the enzymatic pathways and bacterial species responsible for the metabolism of these polyphenols in the intestine are largely unknown.

Various in silico approaches using rule- and/or machine learning-based methods have been developed to predict the metabolism of xenobiotics, e.g., PROXIMAL (Yousofshahi et al., 2015), enviPath (Wicker et al., 2016), Meteor Nexus (Judson et al., 2015), and BioTransformer (Wishart et al., 2022). Other approaches in the field utilize reinforcement learning methods (Yousofshahi et al., 2011), (Delepine et al., 2018), (Koch et al., 2020), (Zheng et al., 2022) to identify novel synthesis pathways in host organisms to produce natural products. However, there have been limited attempts to construct prediction tools for gut microbial metabolism of dietary flavonoids and to investigate the role of enzyme promiscuity. In the present study, we develop a chemical reaction class similarity-based prediction tool for investigating microbial flavonoid metabolism. We analyzed the KEGG database to identify and tabulate Reaction Classes (RClass) that represent the known flavonoid metabolism reactions by plant enzymes. The user can provide an enzyme, a bacterial species or a group of species as inputs. The tool generates predictions for the input’s potential to metabolize dietary flavonoids by comparing the RClasses that are associated with the input enzyme(s) and the key RClasses that are chosen as representatives of gut microbial flavonoid metabolism reactions. Our approach in using RClasses instead of individual reactions allows the tool to introduce a degree of promiscuity instead of relying on matching the target molecule to the exact substrates of known reactions.

We used the above prediction tool to investigate if multiple species can collectively catalyze a multi-step pathway of flavonoid metabolism through promiscuous enzymatic action. We predicted the role of individual gut bacterial species in the degradation of tilianin via *O*-deglycosylation, demethylation and hydrogenation. Using *in vitro* culture experiments, we validated our predictions that *Bifidobacterium longum subsp. animalis*, *Blautia coccoides* and *Flavonifractor plautii* each catalyze particular steps in the pathway and together perform the degradation of tilianin to naringenin. Further, we demonstrated that tilianin and its metabolites have different antioxidant activities in a rat neuronal cell model. Prospectively, the prediction-validation methodology developed in this work could be used to systematically characterize the metabolism of flavonoid metabolites and identify the responsible enzymes and species, directly linking the formation of bioactive flavonoid metabolites to abundance of specific gut bacteria.

## Results

### RClass similarity successfully predicts the activity of known flavonoid-metabolizing plant and bacterial enzymes

To match a given enzyme with its non-native polyphenolic substrates and promiscuous flavonoid metabolism reactions, the tool retrieves the RClasses that are associated with the enzyme(s) from the KEGG database, then finds whether those RClasses are similar to the key RClasses that are chosen as representatives of flavonoid metabolism reactions as detailed in the Methods section. These key flavonoid reactions are *O*-deglycosylation, C-hydrolysis, de/hydrogenation, de/methylation, di/hydroxylation, ring cleavage, anthocyanin/anthocyanidin di/hydroxylation and anthocyanin/anthocyanidin de/hydrogenation reactions. Figure 1 shows the general workflow for the predictions. The output contains the reaction type, similarity score and potential flavonoid substrates.

**Figure 1.**
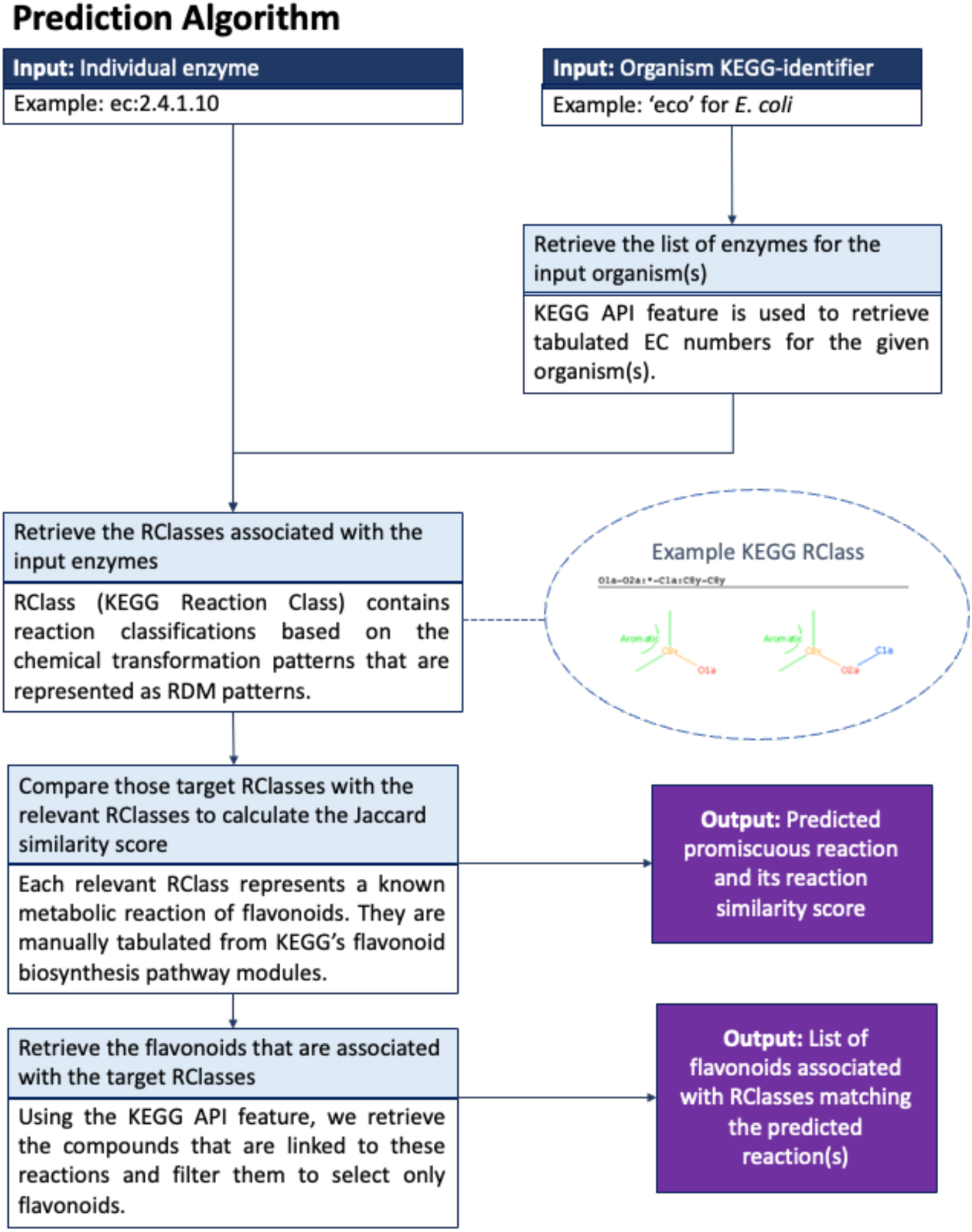
Workflow the algorithm utilizes to generate predictions for a given enzyme or an organism

We evaluated the tool’s accuracy by using known flavonoid-metabolizing enzymes from plants as positive controls, and glycolysis enzymes that are not expected to utilize flavonoids as negative controls (Table 1). The enzymes and their catalytic activities were collected from the BRENDA enzyme database (Schomburg et al., 2004). Positive control enzymes were chosen to test each reaction type that the tool is expected to predict. We could not test the di/hydroxylation of anthocyanin/anthocyanidin as BRENDA does not catalog any enzymes that are known to carry out this reaction. Negative control enzymes were chosen from the Glycolysis/Gluconeogenesis Pathway in the KEGG database to include at least one enzyme from each Enzyme Commission hierarchical group. For the test runs, we included enzymes that catalyze reactions utilizing molecular oxygen, because plant metabolism is not restricted to anaerobic environments.

**Table 1.**
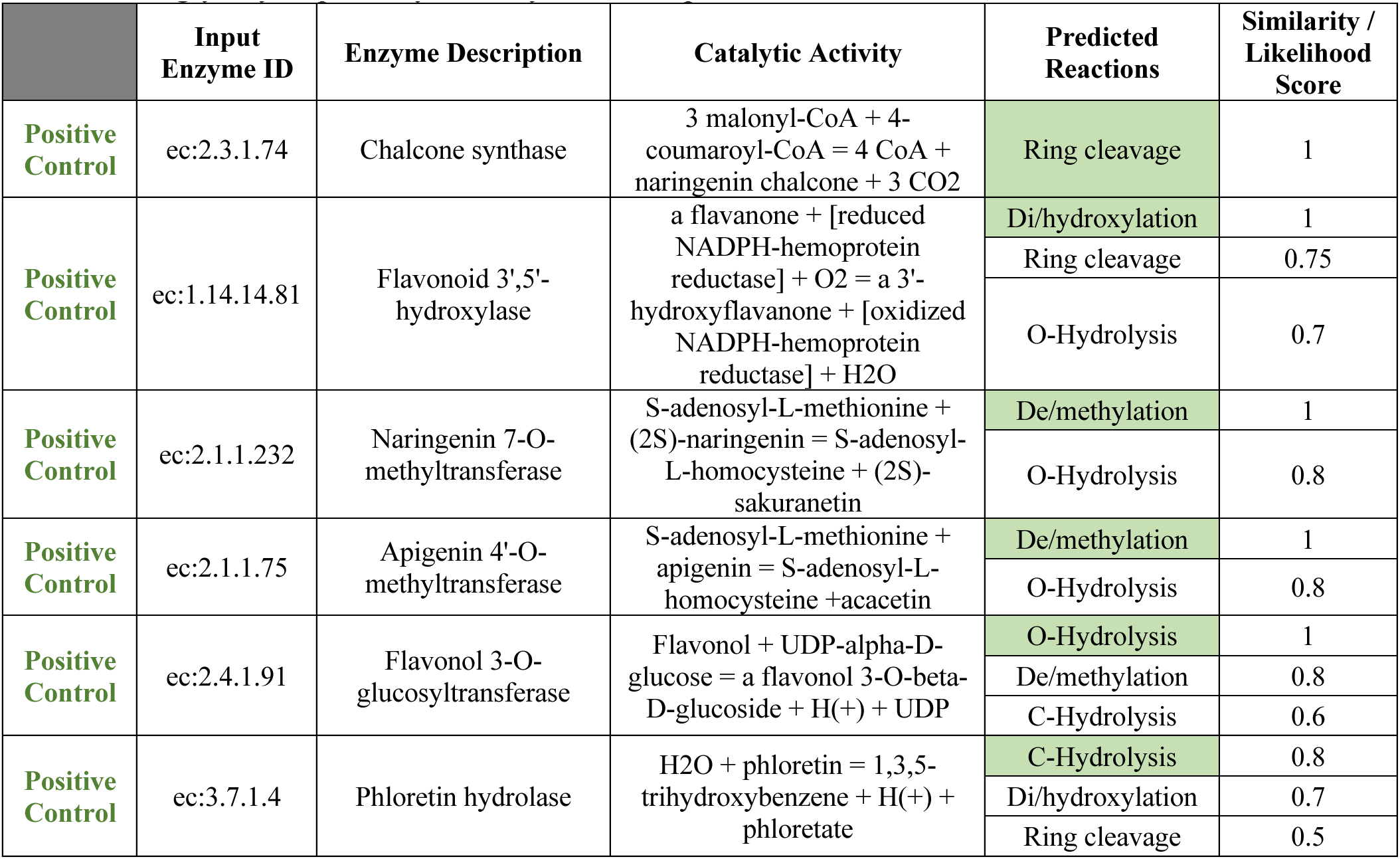
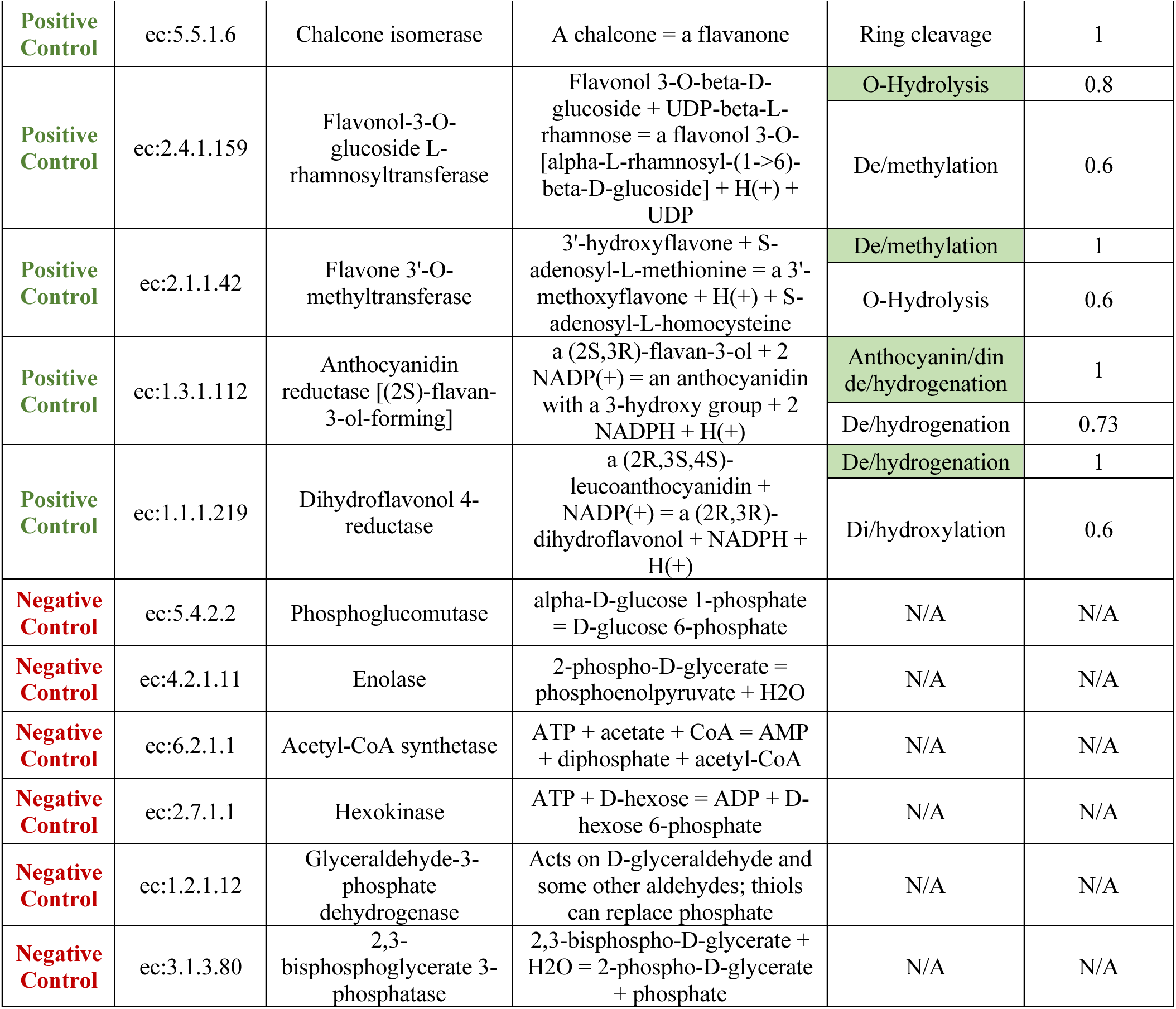
Algorithm testing using positive and negative control examples. Positive controls were chosen among known flavonoid-metabolizing enzymes and negative control enzymes were chosen from glycolysis pathways as they are not expected to metabolize flavonoids.

|  | Input Enzyme ID | Enzyme Description | Catalytic Activity | Predicted Reactions | Similarity / Likelihood Score |
| --- | --- | --- | --- | --- | --- |
| Positive Control | ec:2.3.1.74 | Chalcone synthase | 3 malonyl-CoA + 4-coumaroyl-CoA = 4 CoA + naringenin chalcone + 3 CO<sub>2</sub> | Ring cleavage | 1 |
| Positive Control | ec:1.14.14.81 | Flavonoid 3',5'-hydroxylase | a flavanone + [reduced NADPH-hemoprotein reductase] + O<sub>2</sub> = a 3'-hydroxyflavanone + [oxidized NADPH-hemoprotein reductase] + H<sub>2</sub>O | Di/hydroxylation | 1 |
|  |  |  |  | Ring cleavage | 0.75 |
|  |  |  |  | O-Hydrolysis | 0.7 |
| Positive Control | ec:2.1.1.232 | Naringenin 7-O-methyltransferase | S-adenosyl-L-methionine + (2S)-naringenin = S-adenosyl-L-homocysteine + (2S)-sakuranetin | De/methylation | 1 |
|  |  |  |  | O-Hydrolysis | 0.8 |
| Positive Control | ec:2.1.1.75 | Apigenin 4'-O-methyltransferase | S-adenosyl-L-methionine + apigenin = S-adenosyl-L-homocysteine + acacetin | De/methylation | 1 |
|  |  |  |  | O-Hydrolysis | 0.8 |
| Positive Control | ec:2.4.1.91 | Flavonol 3-O-glucosyltransferase | Flavonol + UDP-alpha-D-glucose = a flavonol 3-O-beta-D-glucoside + H(+) + UDP | O-Hydrolysis | 1 |
|  |  |  |  | De/methylation | 0.8 |
|  |  |  |  | C-Hydrolysis | 0.6 |
| Positive Control | ec:3.7.1.4 | Phloretin hydrolase | H<sub>2</sub>O + phloretin = 1,3,5-trihydroxybenzene + H(+) + phloretate | C-Hydrolysis | 0.8 |
|  |  |  |  | Di/hydroxylation | 0.7 |
|  |  |  |  | Ring cleavage | 0.5 |
| **Positive Control** | ec:5.5.1.6 | Chalcone isomerase | A chalcone = a flavanone | Ring cleavage | 1 |
| **Positive Control** | ec:2.4.1.159 | Flavonol-3-O-glucoside L-rhamnosyltransferase | Flavonol 3-O-beta-D-glucoside + UDP-beta-L-rhamnose = a flavonol 3-O-[alpha-L-rhamnosyl-(1->6)-beta-D-glucoside] + H(+) + UDP | O-Hydrolysis | 0.8 |
|  |  |  |  | De/methylation | 0.6 |
| **Positive Control** | ec:2.1.1.42 | Flavone 3'-O-methyltransferase | 3'-hydroxyflavone + S-adenosyl-L-methionine = a 3'-methoxyflavone + H(+) + S-adenosyl-L-homocysteine | De/methylation | 1 |
|  |  |  |  | O-Hydrolysis | 0.6 |
| **Positive Control** | ec:1.3.1.112 | Anthocyanidin reductase [(2S)-flavan-3-ol-forming] | a (2S,3R)-flavan-3-ol + 2 NADP(+) = an anthocyanidin with a 3-hydroxy group + 2 NADPH + H(+) | Anthocyanin/dinde/hydrogenation | 1 |
|  |  |  |  | De/hydrogenation | 0.73 |
| **Positive Control** | ec:1.1.1.219 | Dihydroflavonol 4-reductase | a (2R,3S,4S)-leucoanthocyanidin + NADP(+) = a (2R,3R)-dihydroflavonol + NADPH + H(+) | De/hydrogenation | 1 |
|  |  |  |  | Di/hydroxylation | 0.6 |
| **Negative Control** | ec:5.4.2.2 | Phosphoglucomutase | alpha-D-glucose 1-phosphate = D-glucose 6-phosphate | N/A | N/A |
| **Negative Control** | ec:4.2.1.11 | Enolase | 2-phospho-D-glycerate = phosphoenolpyruvate + H<sub>2</sub>O | N/A | N/A |
| **Negative Control** | ec:6.2.1.1 | Acetyl-CoA synthetase | ATP + acetate + CoA = AMP + diphosphate + acetyl-CoA | N/A | N/A |
| **Negative Control** | ec:2.7.1.1 | Hexokinase | ATP + D-hexose = ADP + D-hexose 6-phosphate | N/A | N/A |
| **Negative Control** | ec:1.2.1.12 | Glyceraldehyde-3-phosphate dehydrogenase | Acts on D-glyceraldehyde and some other aldehydes; thiols can replace phosphate | N/A | N/A |
| **Negative Control** | ec:3.1.3.80 | 2,3-bisphosphoglycerate 3-phosphatase | 2,3-bisphospho-D-glycerate + H<sub>2</sub>O = 2-phospho-D-glycerate + phosphate | N/A | N/A |

Our analysis shows that Rclass-based similarity can predict the correct reactions for known flavonoid-metabolizing enzymes for ring cleavage, di/hydroxylation, de/methylation, O-deglycosylation, C-hydrolysis, de/hydrogenation and anthocyanin/anthocyanidin de/hydrogenation reactions. We observed that RClass similarity scores for the correctly predicted reactions are above 0.8. Therefore, we set this number as the cutoff value for predictions. Chalcone synthase is classified a transferase enzyme, but we previously found that it can catalyze the C-ring cleavage of naringenin through promiscuous enzymatic action (Gülşan et al., 2022). This was correctly predicted by the tool developed in this work (Table 1). This indicates that our algorithm can also be used to identify enzymes that act on flavonoids to carry out specific metabolism through promiscuous activity. Importantly, our tool did not predict any flavonoid-metabolizing reactions for the glycolysis enzymes as expected.

### Enzyme sequence similarity does not necessarily indicate reaction class similarity

Enzymes having similar protein sequences have been shown to often perform similar catalytic functions. Hence, sequence alignment approaches have been used to classify enzymatic functions. The tool developed in this work instead analyzes functional similarity between enzymes by comparing the enzymes’ reaction classes. To determine if sequence and reaction class similarity are correlated for enzymes involved in *O*-deglycosylation, C-hydrolysis, de/hydrogenation, di/hydroxylation, and de/methylation, we compared the two metrics for these enzymes from five organisms: *E. coli*, *F. plautii*, mouse ear cress, mouse and human. (Table 2)

**Table 2.** Predicted reaction types for the tested organisms mouse ear cress, *F. plautii, E.* coli, human and mouse. Correlation values (R-value) between the sequence similarity and RClass similarity of predicted enzymes, and statistical significance of the correlation value (p-value) have been reported in the last two columns.

| Organism name | Reaction | R-value | p-value |
| --- | --- | --- | --- |
| Mouse Ear Cress | C-hydrolysis | 0.698 | 0.054 |
| Mouse Ear Cress | De/hydrogenation | 0.533 | 0.276 |
| Mouse Ear Cress | O-hydroysis | 0.136 | 0.507 |
| Mouse Ear Cress | Di/hydroxylation | 0.031 | 0.842 |
| Mouse Ear Cress | De/methylation | 0.63 | 0.18 |
| *F. plauti* | Di/hydroxylation | N/A | N/A |
| *F. plauti* | O-hydrolysis | 0.185 | 0.766 |
| *E. coli* | C-hydrolysis | 0.423 | 0.478 |
| *E. coli* | De/hydrogenation | N/A | N/A |
| *E. coli* | De/methylation | 0.866 | 0.333 |
| *E. coli* | Di/hydroxylation | 0.333 | 0.244 |
| *E. coli* | O-hydroysis | 0.567 | 0.018 |
| Human | Di/hydroxylation | 0.563 | 0.245 |
| Human | O-hydroysis | 0.04 | 0.765 |
| Human | De/hydrogenation | 0.086 | 0.711 |
| Mouse | De/hydrogenation | N/A | N/A |
| Mouse | Di/hydroxylation | 0.526 | 0.362 |
| Mouse | O-hydrolysis | 0.327 | 0.011 |

We found no significant correlation between sequence similarity and reaction class similarity in all but one of the aforementioned comparisons. The only exceptions are within the comparison of the enzymes for *O*-deglycosylation in mouse and *O*-hydrolysis in *E. coli*. With a R-value of 0.327, and a p-value of 0.011, there is a weak, albeit significant, correlation in mouse *O*-deglycosylation findings. Similarly, the correlation for *O-*hydrolysis by *E. coli* enzymes show a significant correlation value of 0.567. There are three comparisons (di/hydroxylation in *F. plautii*, de/hydrogenation in *E. coli*, and de/hydrogenation in human) for which neither an R-value nor p-value could be calculated. In all other comparisons, the correlations were insignificant. Four illustrative sample comparisons are shown in Figure 2 for di/hydroxylation in mouse ear cress, de/hydrogenation in human, *O*-deglycosylation in *F. plautii*, and di/hydroxylation in *E. coli*. The correlation is weak in all four comparisons, with R-values of 0.031, 0.086, 0.185, and 0.333, respectively. Furthermore, these correlations are not statistically significant, with p-values of 0.842, 0.711, 0.766, and 0.244, respectively. The result of these comparisons shows that sequence similarity does not necessarily indicate reaction class similarity.

**Figure 2.**
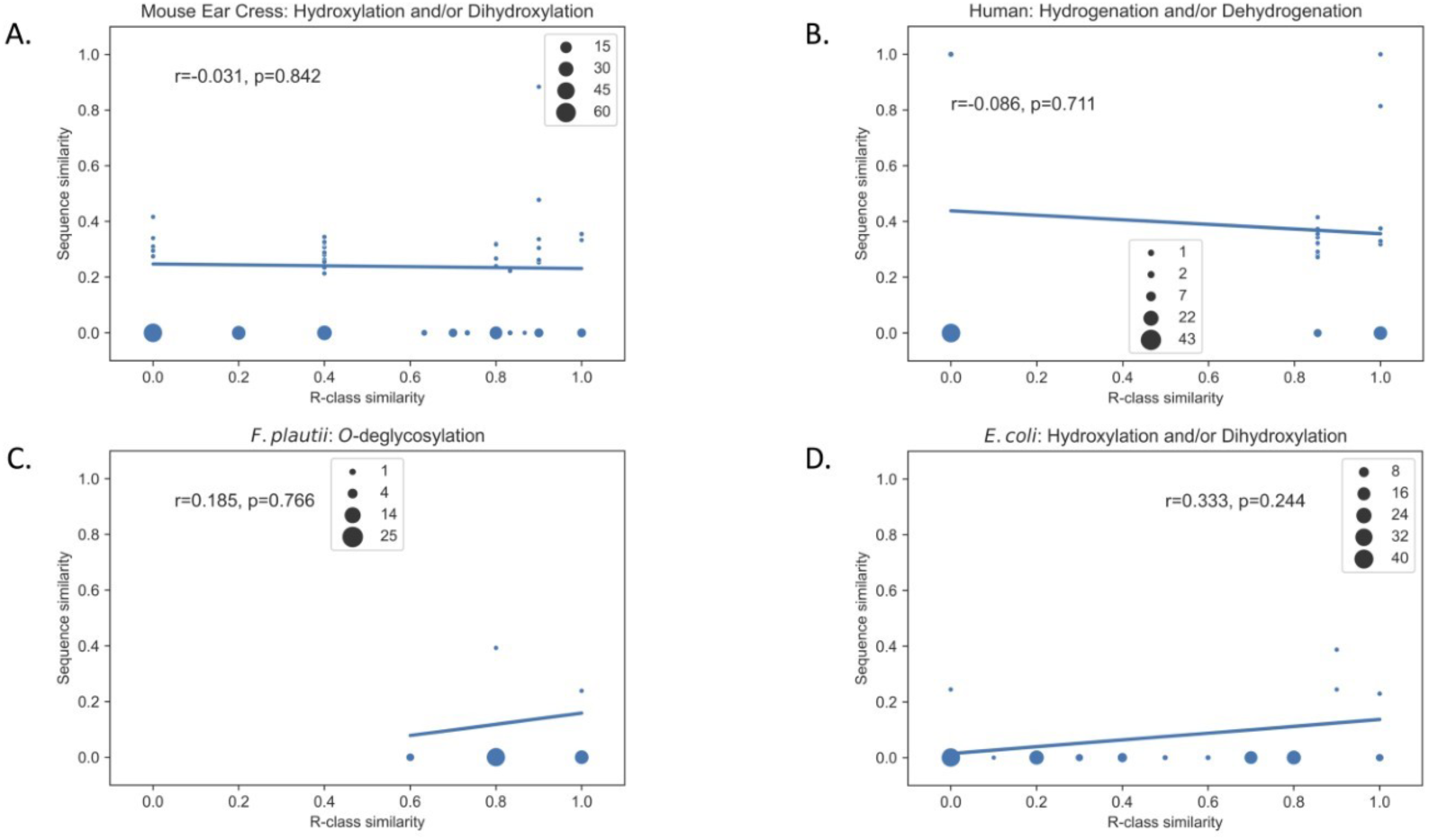
Comparison of sequence similarity and R-class similarity in predicted enzymes for A) di/hydroxylation in mouse ear cress, B) de/hydrogenation in human, C) *O*-deglycosylation of in *F. plautii* and D) di/hydroxylation in *E. coli* The p-values and R-values are calculated in Python using the pearsonr function from the SciPy package.

### Mono- and co-culture experiments confirm tilianin metabolism by selected multiple gut bacteria

We next utilized the prediction tool to investigate the multiple gut bacterial reactions that are required to metabolize a flavonoid glycoside, tilianin, found in many medicinal plants, into an aglycone derivative. Using the tool developed in this work, we are proposing a metabolic pathway that proceeds through *O*-deglycosylation of tilianin to acacetin, then demethylation of acacetin to apigenin, and finally hydrogenation of apigenin to naringenin (Figure 3A). Based on the RClass similarity based predictions, we identified *B. animalis* and *B. coccoides* as gut bacteria capable of the required *O*-deglycosylation and demethylation, respectively. A previous in vitro study identified an ene-reductase enzyme from *F*. *plautii* that can hydrogenate apigenin to naringenin (Schoefer et al., 2004) (Schneider & Blaut, 2000). Thus, we included this reaction in our proposed tilianin metabolism pathway.

**Figure 3.**
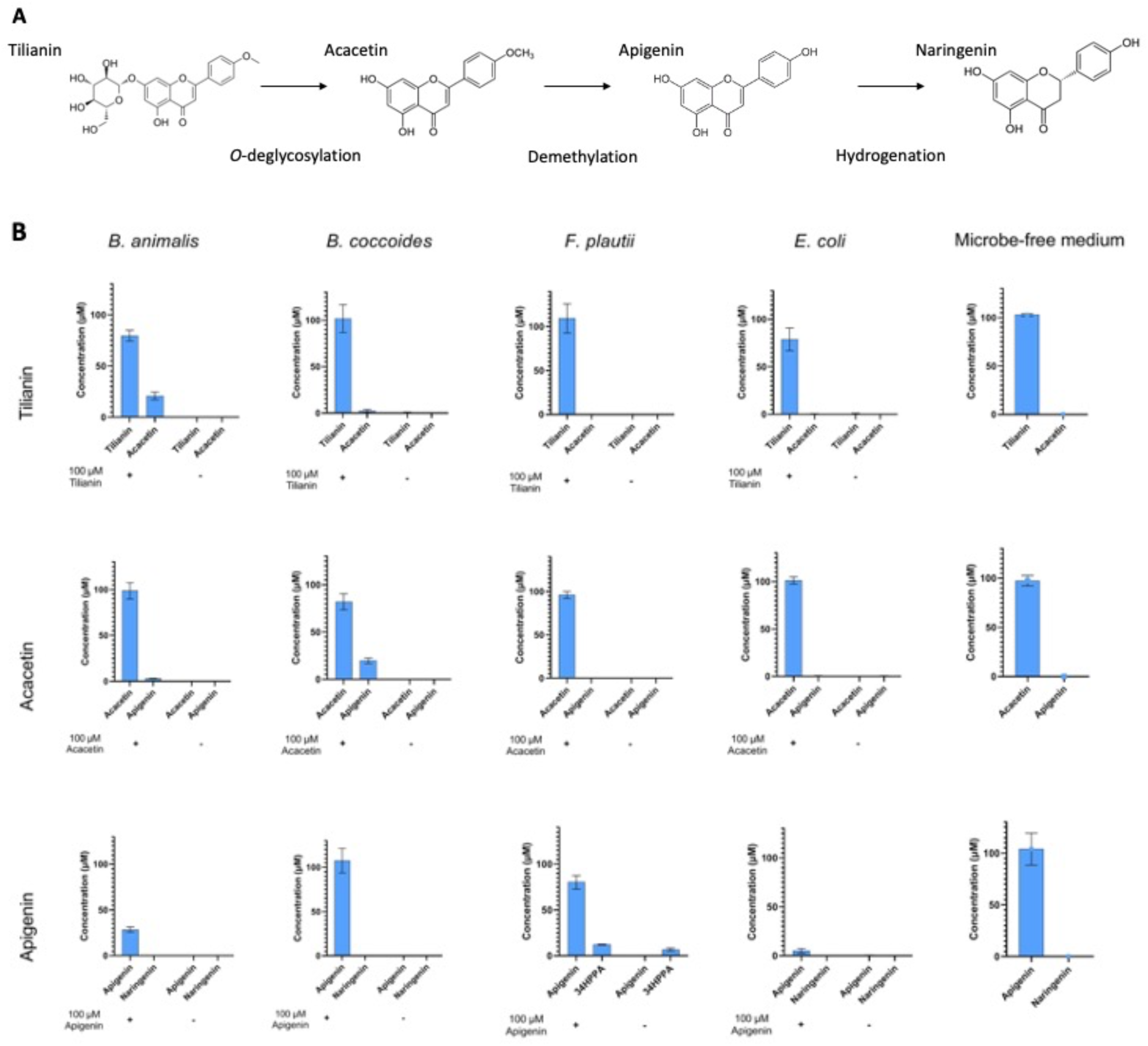
A) Proposed tilianin microbial metabolism pathway. B) Concentrations of parent compounds and their predicted immediate metabolites in the bacterial monocultures of *B. animalis, B. coccoides, F. plautii, E. coli* and microbe-free medium. Results have been represented in bar graphs and displayed together in a matrix format, where the columns represent the microbes, and the rows represent the parent compound that is spiked into the culture.

To experimentally validate the predictions, we cultured selected species anaerobically in the presence of flavonoid substrate of interest for 24 hours, and analyzed the spent medium and cell pellets for accumulation of predicted metabolites. (Figure 3B) We observed that *B. animalis* can convert tilianin to acacetin by removing the glucose group attached to the flavone backbone as predicted. We also observed low concentrations of apigenin in the acacetin treated *B. animalis* culture, suggesting it can also catalyze demethylation on the B-ring of acacetin. While the absence of naringenin in the apigenin treated *B*. *animalis* culture indicates that it cannot hydrogenate apigenin to naringenin, we observed a substantial decrease in apigenin concentration suggesting its consumption by *B. animalis*. We searched for the monoisotopic masses of potential apigenin and naringenin microbial metabolites in tilianin treated *B. animalis* monoculture, which are di/hydroxylation and de/methylation products of apigenin and naringenin. We detected masses for hydroxylated apigenin and methylated naringenin (Supplementary Figure S2). The detected amounts of hydroxylated apigenin and methylated naringenin are statistically significant compared to the vehicle control. We also detected a compound whose mass matches with a dihydroxylated form of apigenin, although the accumulation of this compound was not statistically significant compared to vehicle control. These findings point to further, as yet uncharacterized flavonoid-metabolizing capabilities of *B. animalis*.

*B. coccoides* was predicted to remove the methyl group from acacetin to produce apigenin, but unable to metabolize tilianin and apigenin to acacetin and naringenin, respectively. The in vitro experiments and LC-MS assays confirmed these predictions. Similarly, *F. plautii*, a well-studied flavonoid-metabolizer strain, was not able to utilize tilianin and acacetin. When we incubated *F. plautii* with apigenin, the flavonoid substrate that is shown to be hydrogenated by *F. plautii* (Schoefer et al., 2004) (Schneider & Blaut, 2000), we did not detect any naringenin in the culture despite the decrease in apigenin concentration. However, we observed elevated amounts of 3-(4-hydroxyphenyl)propionic acid (34HPPA), a hydrolysis product of naringenin, in the apigenin treated samples compared to the vehicle control. The concentration difference is statically significant, which points to apigenin hydrogenation to naringenin and its further degradation to 34HPPA. Finally, our predictions also suggested that *E. coli* should not be able to perform any of the proposed tilianin metabolism reactions. Our findings indicate that *E. coli* indeed cannot utilize acacetin over a 24-hour incubation period. Surprisingly, it degraded both tilianin and apigenin in the culture. However, we did not observe any of the predicted metabolites in the E. coli culture, consistent with our predictions that *E. coli* cannot *O*-deglycosylate, demethylate and hydrogenate the selected compounds.

In order to rule out the possibility of chemical degradation of the compounds in the culture medium, we incubated the medium with 100 µM tilianin, acacetin and apigenin and quantified the amount of substrate left in the culture after 24 hours. We did not observe any decrease in the compounds’ concentrations. Further, the predicted degradation products of the selected parent compounds were not present in the culture medium. These results suggest that tilianin, acacetin and apigenin cannot be degraded to their *O*-deglycosylated, demethylated and hydrogenated metabolites in the absence of their metabolizer species.

*O-*deglycosylation of tilianin by *B. animalis* and demethylation of acacetin by *B*. *coccoides* are novel findings that were discovered by using the prediction-validation methodology developed in this work. We next investigated whether these two microorganisms can work together to metabolize tilianin all the way to apigenin. We incubated *B. animalis* and *B*. *coccoides* together and treated the culture medium with 100 µM tilianin for 24 and 48 hours. At the end of 24 hours, roughly 20% of tilianin was converted to acacetin. We observed low concentrations of apigenin, but the amount was not statistically significant compared to the vehicle control from the same timepoint. We observed an increase in acacetin concentration after 48 hours and a significant amount of apigenin formation. (Figure 4A) These results suggest that these two microorganisms can work together to catalyze tilianin *O*-deglycosylation to acacetin and acacetin demethylation to apigenin.

**Figure 4.**
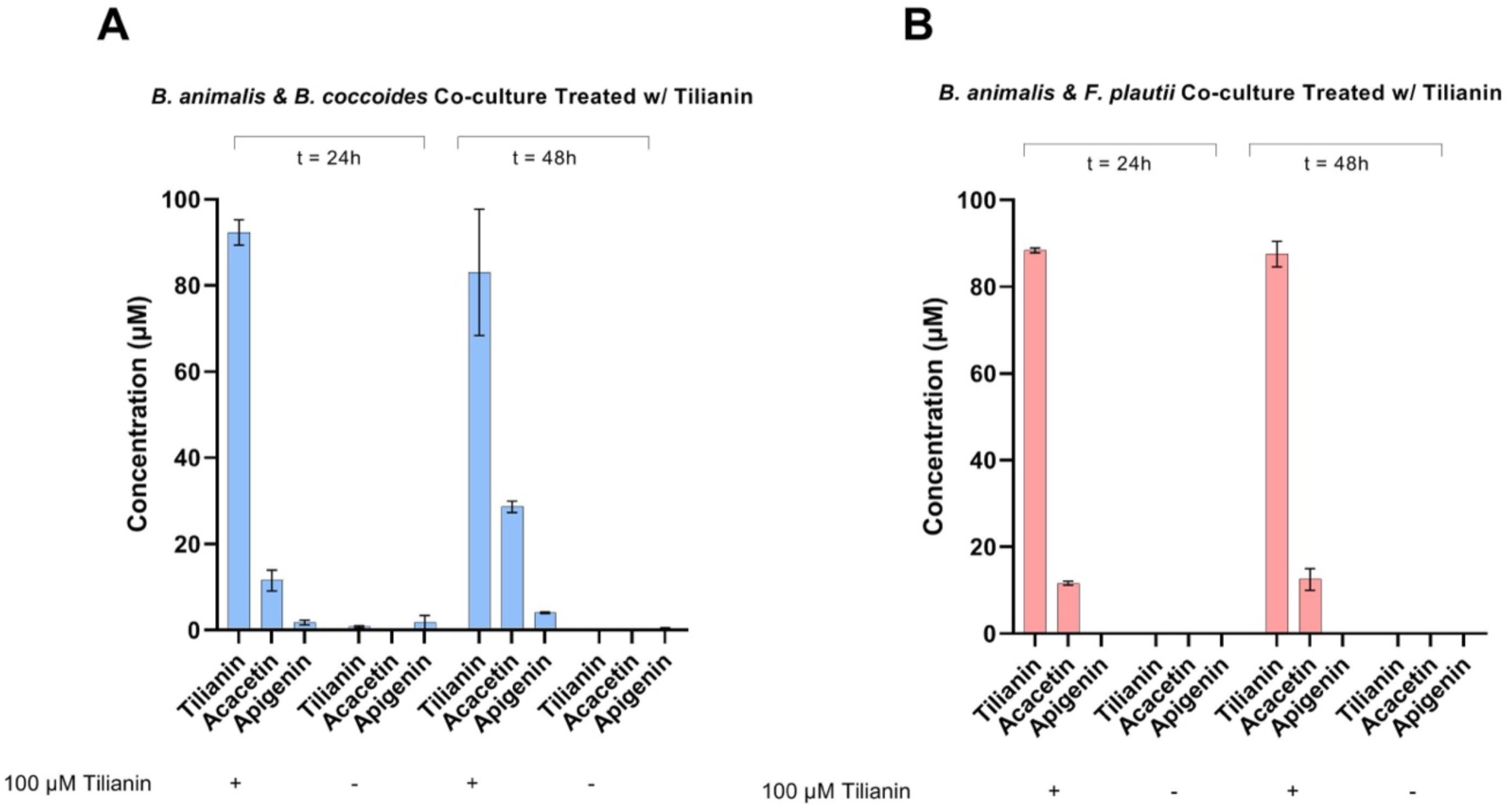
Concentrations of tilianin, acacetin and apigenin in the tilianin-treated co-culture of A*) B. animalis* and *B. coccoides*, and B) *B. animalis* & *F. plautii* after 24 and 48 hours of incubation.

Among the selected bacteria, *B. coccoides* is the only strain that can metabolize acacetin over a 24-hour incubation period (Figure 3). Based on this finding, we hypothesized that this strain is essential for the complete degradation of tilianin. In the absence of *B. coccoides*, acacetin should not be metabolized to apigenin and should accumulate in the culture. We incubated *B. animalis* and *F. plautii* for 24 and 48 hours and treated the culture with 100 µM tilianin. As expected, we did not observe any apigenin formation and acacetin concentration remained stable over time (Figure 4B). This finding indicates that each strain has a unique role in the proposed tilianin metabolism reactions and their unique contribution is essential to carry out the entire pathway.

We next evaluated our predictions against a computational tool that predicts biosynthetic pathways for natural products, BioNavi-NP (Zheng et al., 2022). The analysis was performed by providing tilianin as the query molecule. The tool suggested that the most plausible pathway for tilianin is its *O*-deglycosylation to acacetin, demethylation of acacetin to naringenin, hydrogenation of naringenin to apigenin, and C-ring cleavage to naringenin chalcone (Figure 5). While this pathway overlaps with the predictions from the RClass similarity-based tool developed in this work, BioNavi-NP does not associate these reactions with specific gut bacteria. The enzymes selected by BioNavi-NP are only expressed by plants, whereas our tool links the reactions with gut bacterial enzymes. Another alternative pathway predicted by BioNavi-NP for acacetin degradation to apigenin was through apigenin 7,4-dimethyl ether and genkwanin; however, this pathway is energetically costly and unlikely to occur naturally. BioNavi-NP also suggested that apigenin can be reduced and hydroxylated to 2-hydroxy-2,3-dihydrogenistein, tricetin or 2S-dihydrotricetin. These and other predictions are shown in Figure 5.

**Figure 5.**
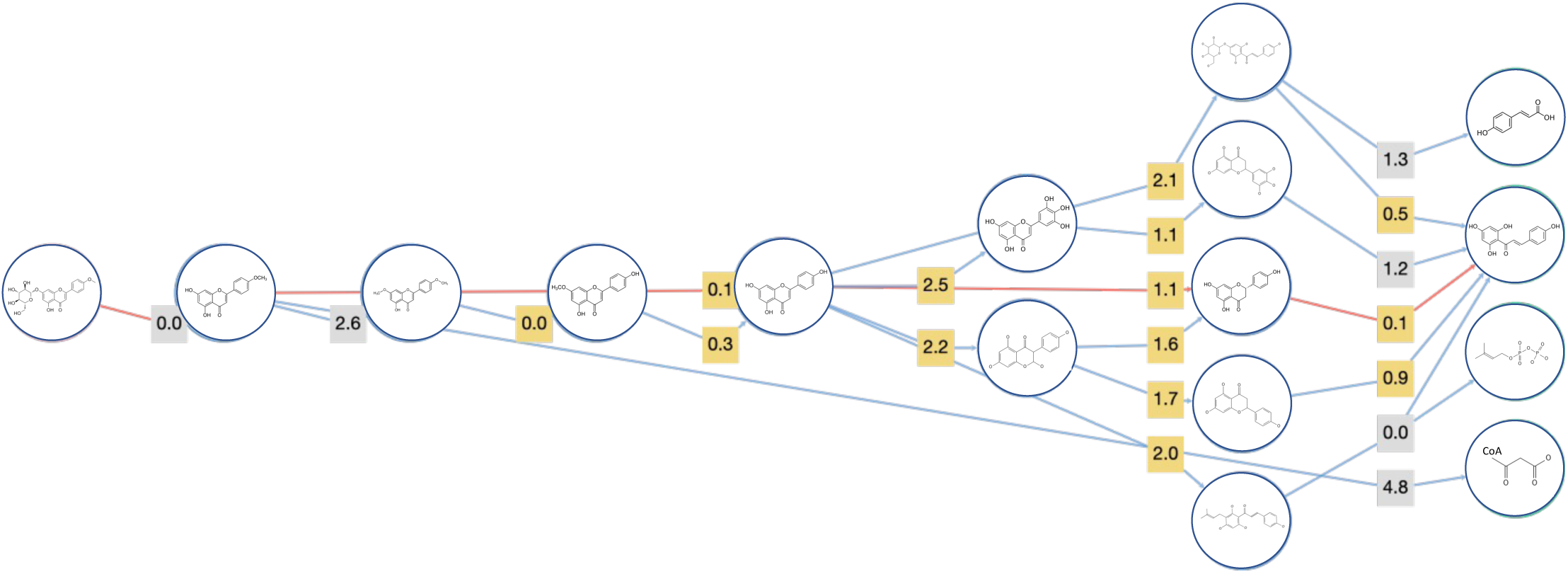
Tilianin metabolism pathway predicted by BioNavi-NP using the algorithm’s default modes. The outputs were redrawn to provide higher resolution for the predicted compounds. The most plausible pathway is colored in red, which overlaps with the pathway proposed in this work. The calculated cost of each reaction step is reflected by the confidence score where lower score means higher reaction probability.

The stoichiometry of tilianin metabolism and conversion yields observed in the mono-culture experiments (Figure 3B) indicate that no other compounds should be formed to any significant extent. Using the monoisotopic masses of the compounds predicted by BioNavi-NP, we performed a mass-based search in our single culture experiments. None of the BioNavi-NP predicted compounds were detected in the cultures (data not shown).

### Tilianin and its microbial metabolites elicit different biological activities

Rat pheochromocytoma cells (PC-12) originally derived from the adrenal gland have been extensively used as in vitro models in neurobiology owing to their ability to depict characteristics similar to cortical neurons (Wiatrak et al., 2020). PC-12 cells are especially desirable for early-stage protective screening against neurodegenerative diseases, which may be mimicked through disease dependent exogenous injury. In this study, we used PC-12 cells for an oxidative damage model as oxidative stress is a prevalent hallmark in several neurological disorders. We investigated the potential of tilianin and tilianin-derived metabolites to protect against hydrogen peroxide (H_2_O_2_) induced oxidative damage to PC-12 cells. Treatment of PC-12 cells with 200 µM H_2_O_2_ sharply decreased cell viability by 67% with an associated increase in ROS level of 57% in comparison to the untreated control group (Figure 3-6). As expected, pretreatment with NAC significantly protected against loss of cell viability and elevation in ROS, with a 43% increase in viability and a 101% reduction in compared to the H_2_O_2_ treated group. Pretreatment with 10 µM acacetin and 5 µM naringenin significantly improved cell viability by 15% and 31% respectively. In contrast, tilianin and apigenin did not significantly impact viability. All tilianin pathway metabolites at 5 µM significantly attenuated H_2_O_2_-induced ROS generation with the greatest reduction of 90% achieved by acacetin. In contrast, tilianin itself failed to reduce ROS generation. Acacetin significantly attenuated ROS also at the higher concentration (10 µM). Except for acacetin, pretreatment with tilianin or any other pathway metabolite at 10 µM or higher concentration did not rescue cell viability or attenuate ROS generation (data not shown).

**Figure 6.**
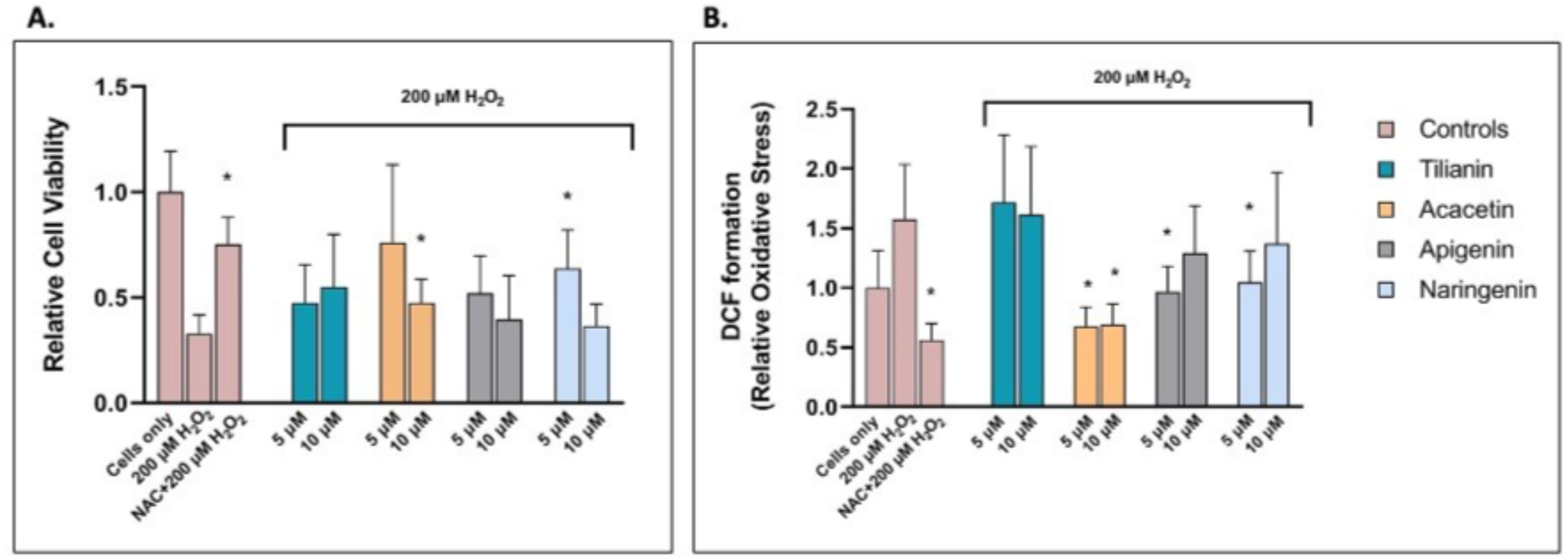
PC-12 response too 200 μM H2O2 stress treatment for 4 hours following 1 hour pretreatment with tilianin, pathway metabolites (acacetin, apigenin, naringenin), or control conditions. A. Viability was detected using calcein violet and shown as fraction of the untreated condition. B. The ROS level was assessed using DCFDA shown as a fraction of the untreated condition. N = 3 (figure shown is representative of all experiments); n = 4; *P < 0.05.

## Discussion

Dietary flavonoids and the commensal gut microorganisms exert a two-way interaction; bacterial metabolism of flavonoids may alter flavonoid bioavailability. Previous studies have repeatedly shown that flavonoid metabolites may elicit different biological activities than their parent compounds, suggesting that gut bacterial species directly impact the profile of bioactive flavonoid metabolites. The most notable examples of microbiome-mediated flavonoid transformation that results in increase in biological activity can be seen in isoflavones (Morishita et al., 1989). Isoflavones are predominantly found in soy and their metabolism is directly linked to the microbiome variability. (Axelson & Setchell, 1981) Recent studies suggest that daidzein conversion to equol takes place in the intestine and is catalyzed by microbial reductase enzymes, however, the key gut microorganisms have not been fully characterized (Mayo et al., 2019). Equol shows stronger estrogenic (Setchell et al., 2005) (Jackson et al., 2011), anti-androgenic (Lund et al., 2004) and antioxidant (Wei et al., 2011) (Wei et al., 2011) activity than its food precursors daidzein and daidzin. Another possible metabolic conversion for daidzein is its C-ring cleavage to O-desmethylangolensin, which is likely produced by different bacterial taxa (Mayo et al., 2019). A majority of humans (80-90%) cannot produce equol as they metabolize a large fraction of daidzein to O-desmethylangolensin, which has no estrogenic activity. (Gardana et al., 2014) (Frankenfeld, 2011) These findings are crucial to understand the gut microbiota-dependent physiological effects of dietary flavonoids. Flavonoids may also alter the composition of microbiota profile by promoting beneficial bacteria as prebiotics, while inhibiting the growth of pathogenic species due to their anti-microbial properties. Studies have shown that flavonoids may stimulate the growth of beneficial bacteria *Lactobacillus spp.* and *Bifidobacterium spp*. in the intestine, while inhibiting the growth of pathogenic bacteria such as *Clostridium spp.* (Jiao et al., 2019) (Saad et al., 2016) (Ponzetto et al., 1988) (Anhe et al., 2015) (Carrera-Quintanar et al., 2018) Thus, flavonoids have the potential to therapeutically target the intestinal microbiome.

Similarly, increasing evidence points to a bidirectional interaction between the gut microbiota and the brain. Microbially mediated metabolites may interact with the central nervous system (CNS) indirectly through signaling pathways often involving the vagus nerve or directly by entering systemic circulation and permeating through the blood brain barrier (Bravo et al., 2011). Alternatively, the CNS impacts gut microbiome function through mediation of the gut environment. In a recent example, a brain injury mouse model noticed compositional changes in caecal microbial population dependent on the extent of brain injury, suggesting a causal link between brain damage and the gut microbiota population diversity (Houlden et al., 2016). As a result, the gut-brain-axis has been implicated as an important modulator of neurodegenerative disease pathology. For example, a study using an Alzhiemer’s disease - like pathology (ADLP^APT^) transgenic mouse model noticed a reduction in neuropathological hallmarks namely, amyloid beta plaque formation, neurofibrillary tangles accumulation, and glial reactivity, with both frequent transfer and transplantation of gut microbiota from healthy mice (Kim et al., 2020). Similar clinical studies point to dysbiosis as a disregulator of the microbial metabolite profile capable of interacting with the CNS (Vogt et al., 2017) (Cattaneo et al., 2017). Despite the growing consensus linking dysbiosis to neurodegenerative diseases, the driving cause of many neuronal diseases still remain unclear, causing difficulty in development of treatment targets. One hypothesis considers oxidative stress resulting from the over accumulation of reactive oxygen species as a significant contributor to neuronal damage. Dysregulated reactive oxygen species (ROS) negatively interact with cellular structures and disrupt critical pathways, which then leads to further disease pathogenesis (Singh et al., 2019). Thus, this study seeks to determine the radical scavenging capacity of microbially mediated metabolites to elucidate the role of microbial flavonoid metabolism on oxidative stress protective potential.

In this work, we present a reaction class-similarity based prediction tool to identify multiple gut bacterial species that can together catalyze a multi-step pathway for flavonoid metabolism through promiscuous enzymatic activity. We predicted and experimentally validated that the sugar moiety attached to the A-ring of tilianin can be removed by *B. animalis* to produce acacetin, and that acacetin can be methylated to apigenin by *B. coccoides*. Recently, Yang et al. (Yang et al., 2021) reported hydrogenation of apigenin to naringenin by *F. plautii*. Based on these findings, we propose a novel gut microbial pathway for tilianin degradation in the intestine. Further, we used a rat neuronal cell model to show that tilianin metabolites can protect against oxidative stress induced loss in cell viability, whereas the parent compound tilianin was ineffective.

Gut microbial metabolism of dietary flavonoids, the largest class of dietary phenolic phytochemicals with over 6,000 structurally distinct compounds (Panche et al., 2016), has garnered wide interest from the scientific community due to the compounds’ health promoting effects. Tilianin is a flavonoid glycoside that can be found in a wide range of medicinal plants (Khattulanuar et al., 2022) with reported benefits of being antioxidant (Nam et al., 2006), anti-inflammatory (Jiang et al., 2019), cardioprotective (Galvez et al., 2015) and neuroprotective (Yang et al., 2021). Despite its reported extensive biological activities and medicinal benefits, the bioactive form of tilianin and its mechanisms of action remain to be elucidated. Tilianin has a flavone backbone with a methyl group attached on its B-ring and a glucose group attached on its A-ring. Because of the functional groups located on its backbone and the double bond on its C-ring, tilianin is prone to *O*-deglycosylation, demethylation and hydrogenation. *O-*deglycosylation followed by demethylation yields apigenin (flavone) by way of acacetin. Hydrogenation of C2-C3 bonds of flavones leads to flavanone formation, which is also the first step for their metabolism through C-ring cleavage. Yang et al. identified a flavone reductase from *F*. *plautii* that catalyzes C-ring hydrogenation of apigenin to form naringenin, a common flavanone (Lagkouvardos et al., 2016). The diversity of metabolic reactions possible for tilianin make it an attractive candidate to study catabolism of dietary flavonoids by gut bacteria. Additionally, tilianin and its metabolites (acacetin, apigenin and naringenin) are all commercially available. Taken together, given its biological activity, chemical structure and commercial availability of metabolic derivatives, tilianin is an attractive model compound to investigate gut microbial metabolism of flavonoid glycosides and investigate whether flavonoid-derived metabolites elicit different physiological activities than their parent compounds.

For many dietary phytochemicals, the metabolic pathways in animals and their gut microbiota are not well characterized. One way to investigate these pathways is to reconstruct heterologous biosynthesis pathways using computational retro-biosynthesis tools (Jeffryes et al., 2018). These tools can mainly be classified into two approaches: knowledge-based approaches (Kuwahara et al., 2016) (Latendresse et al., 2014) that utilize reaction databases such as MetaCyc (Caspi et al., 2020), and rule-based approaches (Hafner et al., 2021) (Finnigan et al., 2021) that match the query molecule to generalized rules such as RetroPath RL (Koch et al., 2020). Knowledge-based tools have limited utility for more complex phytochemicals because reactions responsible for their metabolism are not comprehensively cataloged in databases. Rule-based methods utilize structural knowledge (e.g., subgraph patterns) of the compounds of interest to introduce a degree of generality to the reaction rules and thus are capable of predicting potentially novel metabolic pathways; however, this may lead to invalid predictions of products with structures unlikely found in nature (e.g., epoxides). More recently, tools that use machine learning methods to predict enzymatic reactions have shown the potential for higher performance and more accurate generalizations of reaction rules (Liu et al., 2017).

We evaluated our predictions against a reinforcement learning-based computational tool for predicting natural product biosynthetic pathways, BioNavi-NP. (Zheng et al., 2022) BioNavi-NP uses a tree-based searching algorithm to generate retro-biosynthetic pathways and sorts the predicted reactions based on the computational prediction cost and pathway length. The computational cost reflects the confidence score, or a likelihood of a predicted reaction, where smaller cost means higher reaction probability. The reactions can be further ranked by the taxonomic distance between an optional user-defined organism and the organisms associated with the tool-predicted enzyme. Using default parameters, BioNavi-NP predicted that the most plausible transformation for tilianin is its O-deglycosylation to acacetin, demethylation of acacetin to apigenin, which overlaps with our RClass similarity-based predictions. The outputs from BioNavi-NP also suggested that acacetin can be methylated on its B-ring to apigenin 7,4-dimethyl ether, which can be demethylated on its A-ring to genkwanin. However, these reactions were flagged as likely implausible due to their lower confidence score. The most likely reaction pathway predicted by BioNavi-NP for apigenin is hydrogenation to naringenin followed by ring cleavage of naringenin to naringenin chalcone. The predicted compounds that are marked as “less likely” by BioNavi-NP were not detected in our in vitro metabolomics experiments.

One of the attractive features of BioNavi-NP is that it identifies potential enzymes that can catalyze the predicted reactions. It uses Selenzyme (Carbonell et al., 2018) to rank the pathway output by reaction similarity and taxonomic distance between the organism of candidate enzyme and a user-defined organism. Like our RClass similarity-based analysis, BioNavi-NP also predicted that chalcone synthase can catalyze naringenin conversion to naringenin chalcone. However, it was unable to match the other predicted reactions with specific bacteria or bacterial enzymes. All other matched enzymes from BioNavi-NP’s output belong to known flavonoid pathways in plants. Our RClass similarity-based algorithm not only matches these transformations to known plant enzymes, it can also generate predictions for potential matches with bacterial enzymes.

Another useful feature of our algorithm is that its predictions are linked to functional similarity rather than sequence similarity. As shown in Figure 2, an enzyme’s sequence similarity to another enzyme is not strongly correlated to their reaction class similarity. In this regard, relying on homology searches may limit discovery of enzymes that can catalyze reactions involving non-natural substrates through promiscuous enzyme activity.

Despite the above advantages, the prediction tool developed in this work has several limitations. The algorithm relies on the KEGG database to include the strains and enzymes of interest. For example, Eubacterium *ramulus*, a fully sequenced, known flavonoid-metabolizer strain, does not have a KEGG-identifier. Instead, it is recorded with its taxonomy ID (39490). Only three of its flavonoid relevant enzymes are cataloged: chalcone isomerase (CHI), flavonone/flavonol cleaving reductase (FCR) and phloretin hydrolase (PHY). Hence, a user cannot use this microorganism as an input to test its capability of metabolizing flavonoids through promiscuous enzyme activity. One solution to this problem is to use alternative databases such as the UniProt protein knowledgebase (UniProt, 2021), to manually tabulate the enzymes that can be expressed by the organism of interest, and input this list instead of using the KEGG organism-identifier. We used this alternative approach to predict the reactions that can be catalyzed by *B. coccoides* as it also does not have a KEGG organism-identifier. Further, the flavanone/flavonol cleaving reductase from E. *ramulus* does not have an EC number despite its proven function of hydrogenating the C2-C3 bonds of flavanones and flavonols (Braune et al., 2019). Yang et al. recently discovered a functionally similar enzyme, flavone reductase (FLR), in *F. plautii* (Yang et al., 2021), which can initiate the gut microbial metabolism of flavones and flavonols by cleaving the C2-C3 bonds on the C-ring. Although *F*. *plautii* has a KEGG-identifier (fpla), the KEGG database does not report any homologous ene-reductases that are expressed by *F. plautii*. Hence, our tool failed to predict hydrogenation activity of *F*. *plautii* on flavones when “fpla” was used as input despite the organism’s proven ability to hydrogenate apigenin to naringenin. As FLR also does not have an EC number, we were not able to use this enzyme as a single input. Instead, we had to input enoate reductase (EC: 1.3.1.31), a functionally similar ene-reductase, to predict the final step of the tilianin metabolism pathway.

In the present work, we demonstrated that the proposed tilianin pathway can be catalyzed by the select organisms*: B. animalis, B. coccoides* and *F. plautii*. In a comprehensive study by Lagkouvardos et al. (Lagkouvardos et al., 2016) all the 16s rRNA gene amplicon datasets from stool samples stored in Sequence Read Archive (SRA) were analyzed using the tool Integrative Microbial Next Generation Sequencing (IMNGS) (Leinonen et al., 2011). Their findings indicate that *B. animalis*, *B. coccoides* and *F. plautii* are shared by both human and mouse microbiomes. While *B. animalis* and *B. coccoides* are listed among the dominant strains in both organisms, the abundance of *F*. *plautii* is not very high. (Lagkouvardos et al., 2016)

*O*-deglycosylation of tilianin to acacetin by *B. animalis* and demethylation of acacetin to apigenin by *B*. *coccoides* are novel findings facilitated by the use of the prediction tool developed in this work. Even though these two microbes can catalyze the predicted reactions individually, we cannot conclude that they together can catalyze the series of reactions solely based on the monoculture experiment results. For example, *B*. *animalis* can metabolize tilianin to acacetin, but transport of the metabolite acacetin to the extracellular space is not guaranteed. Further, it is unknown whether the amount of acacetin produced by *B*. *animalis* is enough for *B. coccoides* to utilize it to produce apigenin. In order to address these questions, we incubated the two microbes together and treated them with tilianin. After 48 hours of incubation, we observed a decrease in tilianin concentration and formation of acacetin and apigenin in the culture, indicating that *B*. *animalis* and *B*. *coccoides* together can metabolize tilianin all the way to apigenin.

Redox imbalance causing oxidative stress accumulation has been implicated as an important modulator of disorders involving cognitive decline. Extensive research exists around the use of H_2_O_2_ in in vitro neuronal cell injury models as a precursor to ROS, which invokes cell death through apoptotic processes (Wang et al., 2016). Here, we confirmed that PC-12 treatment with H_2_O_2_ induces cell death, likely through ROS mediated apoptosis. PC-12 cells provide a useful in vitro platform for understanding neurological response because of the extensive characterization of their catecholaminergic activity (Wiatrak et al., 2020). Emerging studies prove the utility of dietary polyphenols, namely flavonoids, as effective scavengers of ROS (Zhang & Tsao, 2016). However, radical scavenging potential may be associated with polyphenol structure and electron donation availability. For example, phenolic acids possessing acetic acid or propenoic acid groups achieved greater antioxidant activity than methoxy or hydroxy groups in the direct scavenging of 2, 2′-diphenyl-1-picrylhydrazyl (DPPH) and reduction of Fe(III) to Fe(II) (Chen et al., 2020). Similarly, flavonoid glycosides and aglycones differentially interact with cellular receptors such as the aryl hydrocarbon receptor (AhR), an important mediator of both prooxidant and antioxidant responses (Vrba et al., 2012). Thus, flavonoid protective activity could also rely on metabolic transformation. The present study determined differential protection against cell death attributed to ROS scavenging capacity of the parent glycoside and its metabolites. Interestingly, we noticed an inverse relationship between treatment dose and protective ability, suggesting potential toxicity and prooxidant behavior in high doses. This trend is consistent with a similar study investigating the protective capacity of luteolin against H_2_O_2_ induced injury of PC-12 cells, which determined concentrations of luteolin greater than 50 µM notably lowered cell viability (McGuigan et al., 1986). In addition to consideration of potential cellular toxicity, the concentrations of tilianin and metabolic pathway metabolites chosen for this study should also consider the concentration range present following human consumption. A study investigating excretion rates of green tea phenolic metabolites measured concentrations ranging from 1-7 µM of epicatechin and epigallocatechin, the primary flavonoids found in green tea, in human urine samples after administration of 1.2 g of green tea solids (Li et al., 2000). Thus, the selection of flavonoid concentrations in the presented study (i.e. 5 µM and 10µM) falls within a realistic range for human metabolism without imposing significant toxicity on the PC-12 test system. Further experiments are warranted to fully characterize the concentration dependent ROS protective effects of phenolic metabolites. Nevertheless, the findings in this work clearly demonstrate a difference in biological activity elicited by a health promoting glycoside and its metabolites. We observed that metabolites that reduced loss of cell viability also reduced the level of ROS, suggesting cell viability is recovered through mitigation of oxidative stress. In contrast, lower viability is associated with increased formation of ROS, implying these metabolites are less effective scavengers.

In summary, this study presents a reaction class similarity-based prediction tool to identify the key taxa required to metabolize dietary flavonoids through investigating promiscuous enzyme activity. We report novel experimental evidence that *B. animalis* can *O-*deglycosylate tilianin to acacetin and *B*. *coccoides* can demethylate acacetin to apigenin, as predicted by the tool developed in this work. Combining our results with the earlier literature findings on apigenin hydrogenation to naringenin by *F. plautii*, we complete the metabolic pathway of tilianin by the gut microorganisms. We further showed that a flavonoid and its metabolites have varying potential to protect against cell death attributed to ROS scavenging capacity as antioxidants using an in vitro rat neuronal cell model. Taken together, these results will be beneficial in understanding the microbial metabolism-dependent biological effects of bioactive dietary flavonoids.

## Experimental Procedures

### Construction of the Prediction Algorithm

This project aims to develop a prediction-validation methodology to identify the key microorganisms that are responsible for flavonoid metabolism in the gut and predict potential promiscuous enzymes that can catalyze these reactions. Our prediction tool can take either a single enzyme in the form of Enzyme Commission (EC) number (e.g. “ec:2.1.1.75”), or a KEGG organism-identifier (e.g. “cpv”) or a consortium, a list of different organism-identifiers, as input. The output file consists of a table with possible reactions the input enzyme(s) or microorganism(s) can catalyze, the flavonoid substrates of these predicted reactions, and reaction likelihood score which is based on RClass similarity calculations. (See the Supplementary Figure S1 for an example input and output)

KEGG (Kyoto Encyclopedia of Genes and Genomes) is a knowledge-based integrated database for systematic analysis of genomic information with higher order functional information. It is classified into four major subcategories, which are Systems Information, Genomic Information, Chemical Information and Health Information. We utilized the Chemical Information subcategory to tabulate the relevant compound, reaction, reaction class (RClass) and enzyme data for our purposes. The tool was constructed in three steps. The first step is to tabulate all flavonoids and their first derivatives from the KEGG database. The second step is to retrieve their associated RClasses to further categorize them into major microbial flavonoid reactions. The last step is to create a master table that lists the calculated RClass similarity scores using the method developed by Muto et al. (Muto et al., 2013)

### Step 1: Tabulating flavonoids from the KEGG database

KEGG PATHWAY is a collection of manually drawn pathway maps representing the knowledge on molecular interactions. We searched map00941 (Flavonoid biosynthesis), map00942 (Anthocyanin biosynthesis), map00943 (Isoflavonoid biosynthesis) and map00944 (Flavone and flavonol biosynthesis) pathways to collect the flavonoids and their first neighbors, as these neighbors represent the closest derivatives of chosen flavonoids. Next, we searched the Phytochemical Compounds list from the KEGG database to retrieve the compounds that are not available in the KEGG PATHWAY collection. We removed aurones, 3-arylcoumarins, pterocarpans, coumestans, malonates, rotenones, 2-aryl benzofurans and complex flavonoids as they do not carry the key flavonoid structure which consists of two phenyl rings and a heterocyclic C-ring containing the embedded oxygen. In the end, we collected 312 unique flavonoids and flavonoid metabolites. (Supplementary Table S1)

### Step 2: Retrieving and classifying the associated RClasses

KEGG RClass contains reaction classifications based on the chemical transformation patterns that are represented as RDM patterns. The RDM patterns are defined based on the reaction center (R), difference region (D) and matched region (M) for each substrate-product pair. RClasses represent each unique RDM pattern or a combination of RDM patterns if more than one reaction center is identified for a chemical transformation. Each RClass is identified by the RC number and represented by KEGG atom types. For example, R09791, methylation reaction of naringenin by naringenin 7-O-methyltransferase, belongs to the RClass RC00392. Its KEGG atom type illustration is O1a-O2a:*-C1a:C8y-C8y, which represents the RDM pattern of this particular reaction. The KEGG atom type usually consists of three characters: the first character, the upper case letter, represents the atomic species (O: oxygen), the second character, the number, represents the primary bonding, and the third character, the lower case letter, indicates the topological information such as the number of substituted group. “*” is used in place of hydrogen atoms.

In this step, we aim to retrieve and classify the RClasses that represent microbial flavonoid reactions that can modify our tabulated 312 unique flavonoids and their derivatives. KEGG API is a REST-style Application Programming Interface to the KEGG database resource. It allows KEGG users to retrieve information for databases of their interest and link related entries by using database cross-references. There is not a direct way to retrieve RClasses from the compound IDs using the KEGG API feature. Thus, one needs to obtain the enzymes that can utilize a compound of interest as substrates first, then link them to their corresponding RClasses. Using this two-step procedure, we first collected the enzymes that are associated with the selected flavonoids and their metabolites in the form of EC numbers. We obtained 100 unique enzymes and added an optional feature to eliminate the ones that catalyze reactions with molecular oxygen, as the majority of the gut microbiome is comprised of obligate anaerobic bacteria (Supplementary Table S2 & S3). Using the tabulated enzymes, we retrieved the corresponding RClasses. We identified 19 unique RClasses that represent the major flavonoid reactions, which are O-deglycosylation, C-hydrolysis, de/hydrogenation, di/hydroxylation, ring cleavage, anthocyanin/anthocyanidin di/hydroxylation and anthocyanin/anthocyanidin de/hydrogenation reactions. We separated the di/hydroxylation and de/hydrogenation reactions for the C-rings of anthocyanins and anthocyanidins, because their reaction centers differ from other flavonoids due to their positive charge. All reactions were considered reversible unless specified otherwise.

### Step 3: Listing the calculated RClass similarity scores

The algorithm predicts promiscuous reactions by comparing the RClasses. As KEGG RClasses are represented using the KEGG atom type format, instead of more conventional identifiers such as InChIKey or SMILES, available chemical similarity clustering cheminformatics toolkits are unable to calculate the similarity scores between different RClasses. Thus, we utilized the RClass chemical similarity calculation method developed by Muto et al. (Muto et al., 2013). Briefly, their method divides RDM representation into binary vectors where each atom type is mapped based on their fingerprints. Fingerprints are represented using letters and they indicate presence or absence of a particular atom, functional groups or bond type. Then those binary vectors are used to calculate the Jaccard similarity coefficient between different RClasses. Calculated scores are normalized to range between 0 and 1. The KEGG database uses this method internally to create a database for pairwise RClass similarity comparisons for each RClass that is listed. We exported those pairwise similarity scores for our 19 unique RClasses that represent the major flavonoid reactions to create a master table (Supplementary Table S4). As a result, once the algorithm takes an enzyme, or a list of enzymes derived from an organism ID (or multiple organism IDs), it retrieves the RClasses that are associated with the input enzymes. Then it finds whether these RClasses are similar to the key RClasses that represent microbial flavonoid metabolism reactions by searching them in the master RClass similarity table. If the similarity score is above the cutoff value of 0.8, the algorithm then retrieves the associated compounds using the KEGG API feature and refine them to only get flavonoids. We chose our cutoff value as 0.8, because when we applied the algorithm to the known flavonoid-metabolizing enzymes, we found that their similarity scores fall above this value as shown in the Results section. Finally, the sorted output contains the reaction type, similarity score and potential flavonoid substrates.

### Prediction of Promiscuous Enzyme Activity of Flavonoid Metabolism

As a proof of concept, we aimed to predict the gut microbial flavonoid reactions and validate these predictions using in vitro culturing methods and targeted metabolomics experiments. We chose tilianin, a flavone glycoside, as the parent compound because it carries a large variety of functional groups and it is commercially available in its pure form along with its proposed metabolites. We proposed a pathway of O-deglycosylation of tilianin to acacetin, aglycone version of tilianin, demethylation of acacetin to apigenin, hydrogenation of apigenin to naringenin and ring cleavage reaction of naringenin to 3-(4-hydroxyphenyl) propionic acid. Naringenin ring cleavage was heavily investigated in a recent study by Gülşan et al. (Gülşan et al., 2022), hence, we excluded this reaction from our analysis. We started our predictions with educated guesses. Bifidobacteriaceae species have been repeatedly reported to remove the sugar moiety from flavonoid glycosides under anaerobic conditions (Marotti et al., 2007)) *B*. *animalis* is a well-studied, fully sequenced, commercially available gut anaerobe, hence, we decided to investigate the predictions on this strain. Our algorithm predicted 38 flavonoid metabolizing reactions for *B*. *animalis*, 30 of them being *O*-deglycosylation reactions. Recent studies report that *B*. *animalis* can catalyze the *O*-deglycosylation of anthocyanidins and isoflavones (Ávila et al., 2009), but its activity on flavones remains to be elucidated. Our algorithm predicts that this strain can perform *O*-deglycosylation reaction on vitexin, prunin, hesperetin 7-*O*-Glucoside, neo-hesperidin and apigetrin. Our predictions are limited to the compounds that are tabulated by the KEGG database. Because tilianin does not have a KEGG identifier, it is expected that our algorithm cannot predict *B. animalis* metabolic activity on tilianin. Apigetrin, on the other hand, is a demethylation product of tilianin whose sugar moiety, A and C rings are identical to tilianin. *B. animalis O*-deglycosylation activity on apigetrin was predicted by our algorithm, which gives us confidence that this strain can also metabolize tilianin to its aglycone form, acacetin.

We followed a similar methodology to make predictions for the demethylation reaction. *Blautia* sp. MRG-PMF1 strain was shown to catalyze demethylation of hesperetin (a flavanone) to eriodictyol and biochanin A (an isoflavone) to genistein (Kim et al., 2014). Similar to *B*. *animalis*, *B. coccoides*, from the same genera with *Blautia* sp. MRG-PMF1, is a fully sequenced common gut anaerobe. Thus, we ran our algorithm using this strain. The algorithm predicts that it can demethylate acacetin to apigenin, thus, we decided to test *B. coccoides* strain for this reaction.

### Tabulating and Comparing Sequence and Functional Similarity Scores of Predicted Enzymes

The sequence similarity of a variety of enzymes from *E coli, F. plautii*, mouse, mouse ear cress, and human are plotted against their functional similarity to demonstrate that sequence similarity does not correlate to functional similarity. The sequence similarities of enzymes were obtained through BLAST^®^ from National Library of Medicine (NIH), and the percent identity in blast result is used as the sequence similarity for the enzymes. DB Search feature of the KEGG database were used to tabulate the functional similarity scores of RClasses that are enabled by the selected enzymes. These sequence similarity scores of each enzyme were then tabulated alongside with the functional similarity scores between all other enzymes of the same reaction class and organism, excluding comparisons with themselves. These data points were then graphically represented by bubble plots. Bubble plots were chosen because there are a significant portion of enzymes that have identical sequence and functional similarity scores. For example, there are 2926 unique enzyme pairing in human *O*-deglycosylation group, but there are only 58 unique sequence and functional similarity score pairing. This calls for another dimension in addition to the x and y axis to showcase the degree of repeats, which is the size of the bubbles (data points). All data points, unique or otherwise, were then used to calculate the r and p values. This was done in Python with the pearsonr function in the SciPy package.

### Cell Lines, Bacterial Strains, Culture Conditions, and Reagents

*B. animalis* ATCC 700541, *B.coccoides* ATCC 29236, *F*. *plautii* ATCC 49531 were obtained from ATCC (Manassas,VA). *B*. *animalis* and *B. coccoides* were cultured in brain heart infusion (BHI) broth (Becton, Dickinson and Company, Franklin Lakes, NJ) supplemented with 0.5% yeast extract (Becton, Dickinson and Company, Franklin Lakes, NJ), 0.05% cysteine (Alfa Aesar, Tewksbury, MA), 0.05% hemin, and 0.02% vitamin K (Sigma-Aldrich, St. Louis MO). F. *plautii* was grown in Wilkens-Chalgren Anaerobe (WCA) Broth (Thermo Fisher Scientific, Waltham, MA). Tilianin, acacetin, apigenin, and naringenin (HPLC grade, ≥95.0% purity) were purchased from Sigma-Aldrich (St. Louis, MO). All other high-purity LCMS-grade reagents were also purchased from Sigma Aldrich unless specified otherwise.

### Bacterial Monoculture and Co-culture

Bacterial monoculture experiments with strict anaerobes *B. animalis*, *B.coccoides*, *F. plautii* and *E. coli* were carried out in an anaerobic chamber with 85% N_2_, 10% CO_2_, 5% H_2_ (Coy Lab, Grass Lake, MI) in their respective media mentioned above. All reagents and media were introduced into the anaerobic chamber a day before the experiment to ensure that they were pre-reduced by the start of the experiment. Cultures grown to an OD (absorbance at 600 nm) of 1.0 were diluted 10-fold for *B. animalis* and *B*. *coccoides* and treated with 100 μM of tilianin and acacetin, respectively (stock solutions were prepared using DMSO as solvent and then filter sterilized). *F*. *plautii* monoculture was treated with 100 μM apigenin without any dilution due to its faint growth. Monocultures treated with 0.1% v/v DMSO were used as vehicle controls. Following the flavonoid or vehicle treatment, monocultures were incubated at 37 °C for 48 h. Each monoculture experiment was run in triplicate. Samples were collected for metabolite extraction by sacrificing the cultures after 24 h of incubation.

For bacterial co-culture, *B. animalis* and *B. coccoides* cultures grown to an OD_600_ of 1.0 were pelleted down by centrifugation at 10,000g for 15 min (Eppendorf 5424R) and resuspended in 2.5mL of BHI each and then put together (final culture volume is 5 mL). This co-culture sample was then treated with 100 μM of tilianin and incubated at 37 °C for next 48 h. Each co-culture experiment was run in triplicates under anaerobiosis. Samples were collected for metabolite extraction by sacrificing the cultures after 24 and 48 h of incubation.

For the co-culture of *B. animalis* and *F. plautii*, monocultures grown to an OD_600_ of 0.2 were pelleted down by centrifugation at 10,000g for 15 min (Eppendorf 5424R) and resuspended in 2.5mL of BHI each and then put together (final culture volume is 5 mL). The OD was comparatively lower due to the faint growth of *F. plautii* even after 24h of culture. This co-culture samples were then treated with 100 μM of tilianin and incubated at 37 °C for next 48 h. Each co-culture experiment was run in triplicates under anaerobiosis. Co-cultures treated with 0.1% v/v DMSO and maintained in the same conditions were used as vehicle controls. Both treatment and control samples were collected for metabolite extraction by sacrificing the cultures after 24 and 48 h of incubation.

### Metabolite Extraction

Metabolites were extracted from the harvested cell culture samples using a solvent-based method. Briefly, 50 µL of culture sample was mixed with 200 µL of ice-cold methanol using a vortex mixer and kept on ice for 60 minutes. The sample-solvent mixture was then centrifuged at 15,000g at 4 °C for 5 min. The supernatant was filtered through a 0.22 µm sterile nylon filter (Corning). The filtrate was then passed through a 3 kDa molecular weight cutoff filter (Amicon Ultra, Sigma-Aldrich) placed in a collection tube by centrifuging the sample tube at 15,000g for 45 min at 4 °C. The filtrate from the collection tube was then mixed with 2 mL sterile water, lyophilized to a pellet using a freeze dryer (FreeZone, Labconco Corporation) and stored at -80 °C until further analysis. Prior to LC-MS/MS analysis, the freeze-dried samples were reconstituted in 250 µL of 50% (v/v) methanol/water.

### Targeted Analysis of Tilianin Metabolites

The extracted samples were analyzed for tilianin, acacetin, apigenin and naringenin using product ion scan experiments performed on a triple-quadrupole time-of-flight instrument (5600+, AB Sciex) coupled to a binary pump high-performance liquid chromatography (HPLC) system (1260 Infinity, Agilent). Tilianin was detected in positive ionization mode, while its metabolites were detected using negative ionization mode. Chromatographic separation was achieved on a Synergi 4 µm Hydro, 250 mm × 2 mm, 80 A reverse phase column (Phenomenex, Torrance, CA) maintained at 30 °C using a solvent gradient method. Solvent A was 0.2% (v/v) formic acid in water and solvent B was 100% acetonitrile. The gradient method was as follows: 0–30 min (95% A to 45% A), 30–38 min (45% A to 95% A). The flow rate was set to 0.6 mL/min. The injection volume was 10 µL. Raw data were processed in PeakView 2.1 to determine the ion peaks. The peak areas in the extracted ion chromatograms were manually integrated using MultiQuant 2.1 (AB Sciex). The identities of detected compounds were confirmed by matching their retention time and/or MS/MS spectra to high-purity standards run on the same instrument using the same method. De/methylated and di/hydroxylated apigenin and naringenin metabolites and the compounds predicted by BioNavi-NP were only searched by their monoisotopic masses in time-of-flight experiments and were not confirmed with their high-purity standards.

### Investigation of in vitro Protective Effect of Flavonoids in a Neuronal Cell Model

#### Cell Culture

Rat adrenal pheochromocytoma PC-12 cell lines were obtained from ATCC (CRL-1721). Cells were cultured at 37 °C at 5% CO_2_. Cells were cultured in Rosewell Park Memorial Institute 1640 medium (RPMI, ThermoFisher Scientific, Waltham, MA) supplemented with 10% horse serum (ThermoFisher Scientific, Waltham, MA), 5% FBS (ATCC, Manassas, VA), 50 U/mL Penicillin-streptomycin (ThermoFisher Scientific, Waltham, MA), and 2 g/L of NaHCO_3_(Sigma-Aldrich, St. Louis, MO). Two thirds of the medium was exchanged every other day for fresh medium and cells were passaged approximately every 4 days. All well plates and culture flasks were coated prior to seeding with 7 µg/mL rat tail tendon collagen I (Sigma-Aldrich, St. Louis, MO) and rinsed with PBS 3 times. PC-12 cells used for experiments were in the exponential growth phase of cell growth between passage 4-6.

PC-12 cells were seeded into 96-well plates coated with collagen I (ThermoFisher, Waltham, MA) and rinsed with 3 volumes of PBS. The cells were seeded at a density of 5 x 10^5^ cells/mL then allowed to adhere for 24 hours. PC-12 cells were then cultured in fresh medium containing 0 µM, 100 µM, 200 µM, 500 µM H_2_O_2_ (Sigma-Aldrich, St. Louis, MO) for 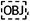4 hours to determine the optimal stress concentration for subsequent experiments (data not shown). Based on cell survival rate, 200 µM H_2_O_2_ was chosen as the oxidative stress concentration.

#### PC-12 Viability Assay

PC-12 cells were seeded in collagen I coated 96-well plates at 5 x 10^5^ cells/mL and allowed to adhere for 24 hours. Cells were then pretreated with fresh medium containing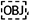 5 µM or 10 µM of either tilianin or tilianin pathway metabolite in quadruplicate. The control conditions received only fresh medium without pretreatment metabolite or 1mM N-acety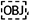l cysteine (NAC, Sigma-Aldrich, St. Louis, MO) as the positive control. Cells were pretreated for 1 hour then exposed to 200 µM H_2_O_2_ for 4 hours. Cell viability was determined using the CellTrace^TM^ Calcein violet live cell stain (ThermoFisher, Waltham, MA). The cell membrane permeable non-fluorescent calcein AM is cleaved by nonspecific intracellular esterase to form highly fluorescent calcein detected at an excitation wavelength and emission wavelength of 400 nm and 452 nm respectively. Cells were first washed twice with PBS then stained with 5 µM calcein in the dark for 1 hour. Fluorescence measurements were then taken using a SpectraMax M3 Multi-Mode Microplate Reader (Molecular Devices, San Jose, CA).

#### Detection of Reactive Oxygen Species

The 2,7-dichlorodihydrofluroescin diacetate (H_2_DCFDA, Sigma-Aldrich, St. Louis, MO) probe was used to measure reactive oxygen species. The cell permeable nonfluorescent H_2_DCFDA is cleaved by intracellular esterase and oxidized to highly fluorescent dichlorofluorescein (DCF) detected with an excitation wavelength and emission wavelength of 492 nm and 517 nm respectively. The measurement of reactive oxygen species followed the same procedure used to determine cell viability. Briefly, PC-12 cells were seeded into 96-well plates and cultured for 24 hours for adherence to the collagen I coating. Cells were then pretreated with 5 µM or 10 µM of either tilianin or tilianin pathway metabolite for 1 hour in quadruplicate. Experimental controls received only fresh medium or pretreatment with 1mM NAC for 1 hour. The cells were then stressed with 200 uM H_2_O_2_ for 4 hours. Treated cells were washed twice with PBS and stained with 20 µM H_2_DCFDA in the dark for 1 hour. Fluorescence was measured using the 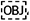 SpectraMax M3 Multi-Mode Microplate Reader (Molecular Devices, San Jose, CA).

## Supporting information

Supplementary Figure S1. Example input and output

Supplementary Table S1. Tabulated flavonoids and their derivatives

Supplementary Table S2 - Tabulated enzymes from the chosen flavonoids and their derivatives

Supplementary Table S3 - Enzymes that catalyze reactions which require oxygen

Supplementary Table S4 - Master RClass similarity table with a cutoff value of 0.8

Supplementary Table S5 - Tool predictions for possible promiscuous enzymes for the proposed Tilianin pathway

Supplementary Figure S2. Concentrations of possible microbial metabolites of apigenin and naringenin detected in the apigenin treated B. animalis mono

## Acknowledgements

This work was in part supported by NIH award #AT010282 (to A.J. and K.L). The authors acknowledge the use of Integrated Metabolomics Analysis Core (IMAC) at Texas A&M University and Mass Spectrometry Core Lab at Tufts University.

---

**Figure S1.**
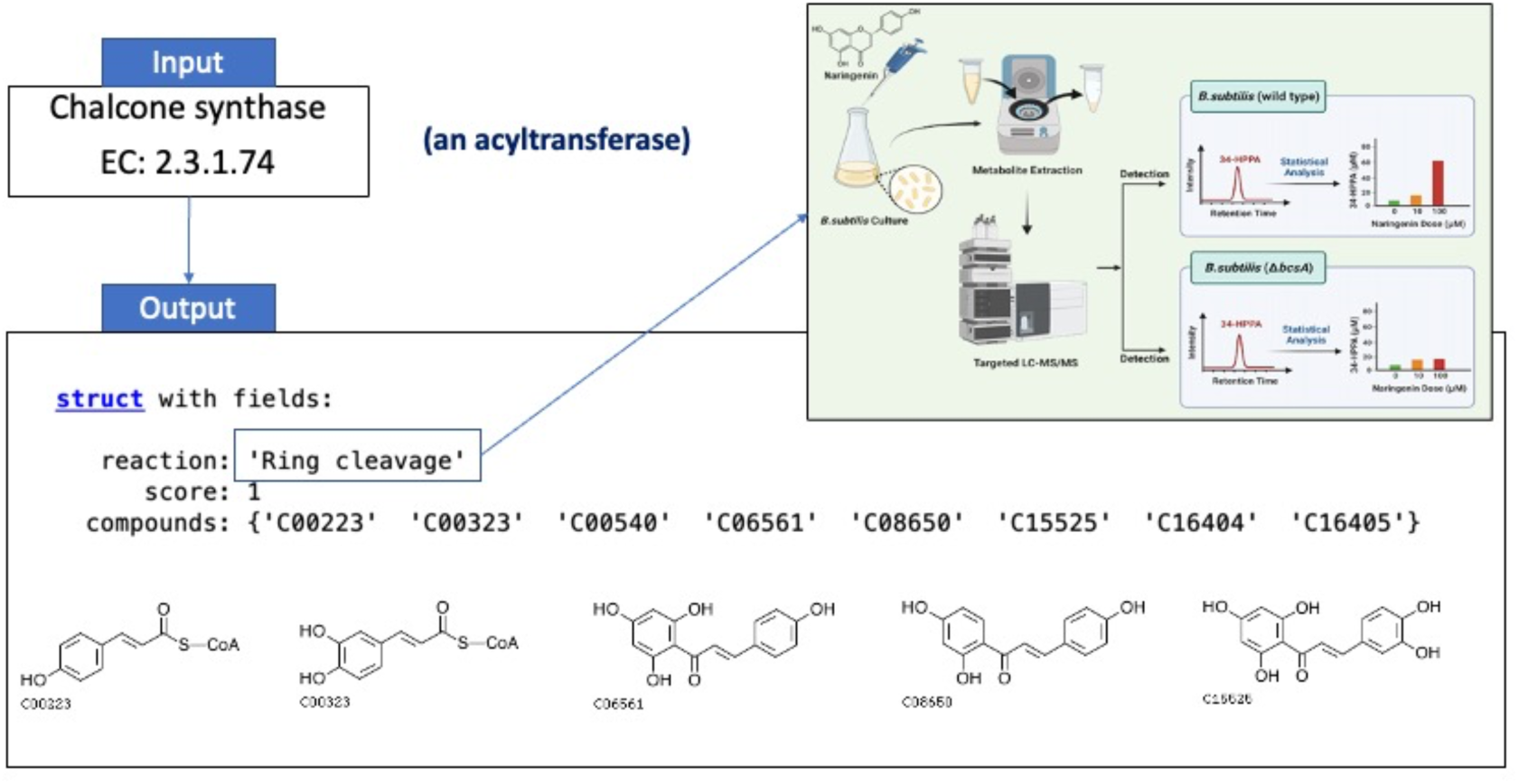
Example input and output from the tool developed in this work We used chalcone synthase enzyme with the EC number 2.3.1.74 as an input. The output suggests that this input enzyme can perform ring cleavage reaction on flavonoids and their metabolites, which aligns with our findings from Chapter 2.

**Figure S2.**
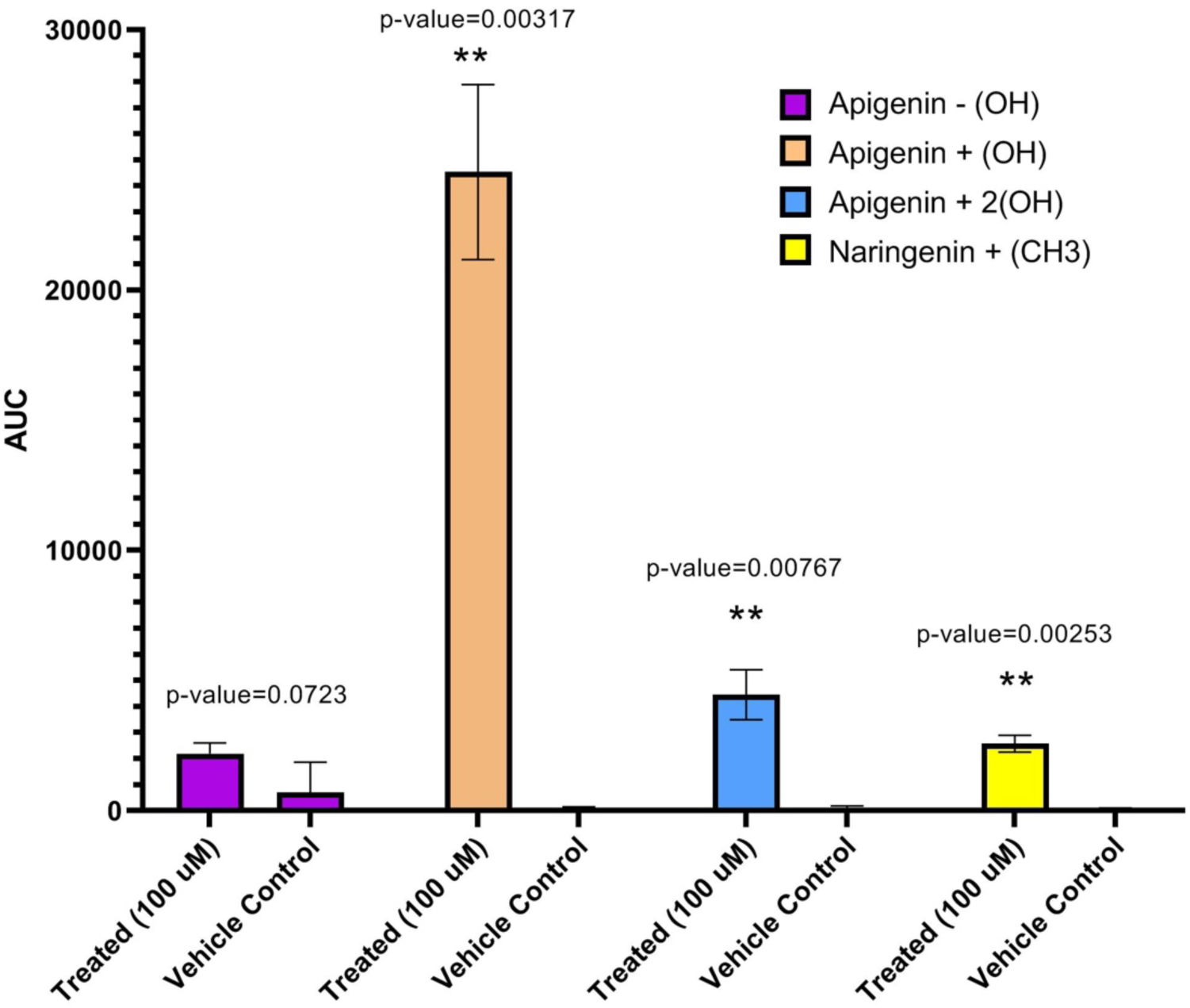
Concentrations of possible microbial metabolites of apigenin and naringenin detected in the apigenin treated *B. animalis* monoculture. Area under the curve for the detected apigenin and naringenin metabolites. Double sterisk (**) indicates significant difference (p < 0.005) compared to the vehicle control (no apigenin).

**Table S1.** Tabulated flavonoids and their derivatives (KEGG COMPOUND numbers)

|  |  |  |  |  |  |
| --- | --- | --- | --- | --- | --- |
| C00223 | C05903 | C09755 | C10097 | C10507 | C14313 |
| C00323 | C05904 | C09756 | C10098 | C10509 | C14314 |
| C00389 | C05905 | C09757 | C10102 | C10510 | C14344 |
| C00509 | C05906 | C09762 | C10104 | C10513 | C14472 |
| C00540 | C05907 | C09770 | C10105 | C10520 | C14536 |
| C00774 | C05908 | C09774 | C10107 | C10521 | C14942 |
| C00786 | C05909 | C09789 | C10108 | C10524 | C14953 |
| C00814 | C05911 | C09793 | C10109 | C10526 | C14985 |
| C00852 | C05990 | C09803 | C10110 | C10527 | C15019 |
| C00858 | C06561 | C09805 | C10111 | C10530 | C15109 |
| C00974 | C06562 | C09806 | C10112 | C10533 | C15118 |
| C01265 | C06563 | C09807 | C10113 | C10534 | C15509 |
| C01378 | C08064 | C09826 | C10114 | C10536 | C15525 |
| C01460 | C08574 | C09827 | C10115 | C10538 | C15531 |
| C01470 | C08578 | C09828 | C10116 | C10540 | C15567 |
| C01477 | C08600 | C09830 | C10117 | C11264 | C15608 |
| C01514 | C08604 | C09833 | C10118 | C11577 | C16187 |
| C01562 | C08620 | C09939 | C10119 | C11620 | C16188 |
| C01604 | C08629 | C09982 | C10120 | C11621 | C16190 |
| C01617 | C08639 | C10019 | C10121 | C12123 | C16195 |
| C01709 | C08647 | C10021 | C10122 | C12124 | C16225 |
| C01714 | C08649 | C10022 | C10175 | C12125 | C16232 |
| C01750 | C08650 | C10023 | C10176 | C12127 | C16233 |
| C01821 | C08652 | C10024 | C10177 | C12128 | C16298 |
| C02495 | C08716 | C10025 | C10178 | C12134 | C16305 |
| C02549 | C08718 | C10026 | C10180 | C12135 | C16306 |
| C02675 | C08724 | C10028 | C10181 | C12136 | C16307 |
| C02806 | C08725 | C10029 | C10183 | C12137 | C16312 |
| C02906 | C08729 | C10030 | C10184 | C12138 | C16314 |
| C02920 | C09099 | C10031 | C10185 | C12139 | C16315 |
| C03515 | C09126 | C10032 | C10186 | C12140 | C16404 |
| C03567 | C09318 | C10033 | C10187 | C12208 | C16405 |
| C03648 | C09320 | C10036 | C10188 | C12249 | C16406 |
| C03870 | C09471 | C10037 | C10189 | C12625 | C16407 |
| C03951 | C09478 | C10038 | C10190 | C12626 | C16408 |
| C04024 | C09505 | C10039 | C10191 | C12627 | C16409 |
| C04109 | C09614 | C10040 | C10192 | C12628 | C16410 |
| C04293 | C09616 | C10041 | C10193 | C12629 | C16415 |
| C04443 | C09617 | C10043 | C10195 | C12630 | C16416 |
| C04444 | C09618 | C10044 | C10196 | C12631 | C16417 |
| C04491 | C09636 | C10045 | C10197 | C12633 | C16418 |
| C04527 | C09641 | C10046 | C10199 | C12634 | C16419 |
| C04552 | C09644 | C10047 | C10200 | C12635 | C16422 |
| C04581 | C09646 | C10049 | C10203 | C12636 | C16490 |
| C04608 | C09650 | C10050 | C10205 | C12638 | C16492 |
| C05334 | C09727 | C10058 | C10208 | C12639 | C16911 |
| C05376 | C09728 | C10068 | C10216 | C12644 | C19727 |
| C05622 | C09732 | C10072 | C10415 | C12667 | C19796 |
| C05623 | C09734 | C10073 | C10419 | C12668 | C21833 |
| C05625 | C09735 | C10078 | C10420 | C14079 | C21854 |
| C05631 | C09736 | C10079 | C10465 | C14131 | C21913 |
| C05902 | C09751 | C10084 | C10471 | C14290 | C22048 |

**Table S2.** Tabulated enzymes from the chosen flavonoids and their derivatives

|  |  |  |  |
| --- | --- | --- | --- |
| ec:1.1.1.219 | ec:1.3.1.77 | ec:2.3.1.116 | ec:2.4.1.254 |
| ec:1.1.1.234 | ec:2.1.1.104 | ec:2.3.1.133 | ec:2.4.1.286 |
| ec:1.1.1.348 | ec:2.1.1.150 | ec:2.3.1.153 | ec:2.4.1.294 |
| ec:1.14.11.9 | ec:2.1.1.153 | ec:2.3.1.170 | ec:2.4.1.297 |
| ec:1.14.14.135 | ec:2.1.1.155 | ec:2.3.1.171 | ec:2.4.1.298 |
| ec:1.14.14.81 | ec:2.1.1.169 | ec:2.3.1.173 | ec:2.4.1.299 |
| ec:1.14.14.82 | ec:2.1.1.175 | ec:2.3.1.215 | ec:2.4.1.300 |
| ec:1.14.14.87 | ec:2.1.1.212 | ec:2.3.1.74 | ec:2.4.1.357 |
| ec:1.14.14.88 | ec:2.1.1.231 | ec:2.4.1.105 | ec:2.4.1.81 |
| ec:1.14.14.89 | ec:2.1.1.232 | ec:2.4.1.106 | ec:2.4.1.91 |
| ec:1.14.14.90 | ec:2.1.1.254 | ec:2.4.1.115 | ec:2.4.2.25 |
| ec:1.14.14.91 | ec:2.1.1.267 | ec:2.4.1.116 | ec:2.4.2.35 |
| ec:1.14.14.93 | ec:2.1.1.270 | ec:2.4.1.159 | ec:2.4.2.50 |
| ec:1.14.14.96 | ec:2.1.1.338 | ec:2.4.1.170 | ec:2.4.2.51 |
| ec:1.14.19.63 | ec:2.1.1.339 | ec:2.4.1.185 | ec:2.5.1.36 |
| ec:1.14.19.76 | ec:2.1.1.42 | ec:2.4.1.189 | ec:2.8.2.25 |
| ec:1.14.20.4 | ec:2.1.1.46 | ec:2.4.1.190 | ec:2.8.2.26 |
| ec:1.14.20.5 | ec:2.1.1.75 | ec:2.4.1.191 | ec:2.8.2.27 |
| ec:1.14.20.6 | ec:2.1.1.76 | ec:2.4.1.234 | ec:2.8.2.28 |
| ec:1.17.1.3 | ec:2.1.1.78 | ec:2.4.1.236 | ec:3.2.1.167 |
| ec:1.21.3.6 | ec:2.1.1.82 | ec:2.4.1.237 | ec:3.2.1.168 |
| ec:1.3.1.117 | ec:2.1.1.83 | ec:2.4.1.238 | ec:3.2.1.31 |
| ec:1.3.1.45 | ec:2.1.1.84 | ec:2.4.1.239 | ec:4.2.1.105 |
| ec:1.3.1.46 | ec:2.1.1.88 | ec:2.4.1.240 | ec:4.2.1.139 |
| ec:1.3.1.51 | ec:2.3.1.115 | ec:2.4.1.253 | ec:5.5.1.6 |

**Table S3.**
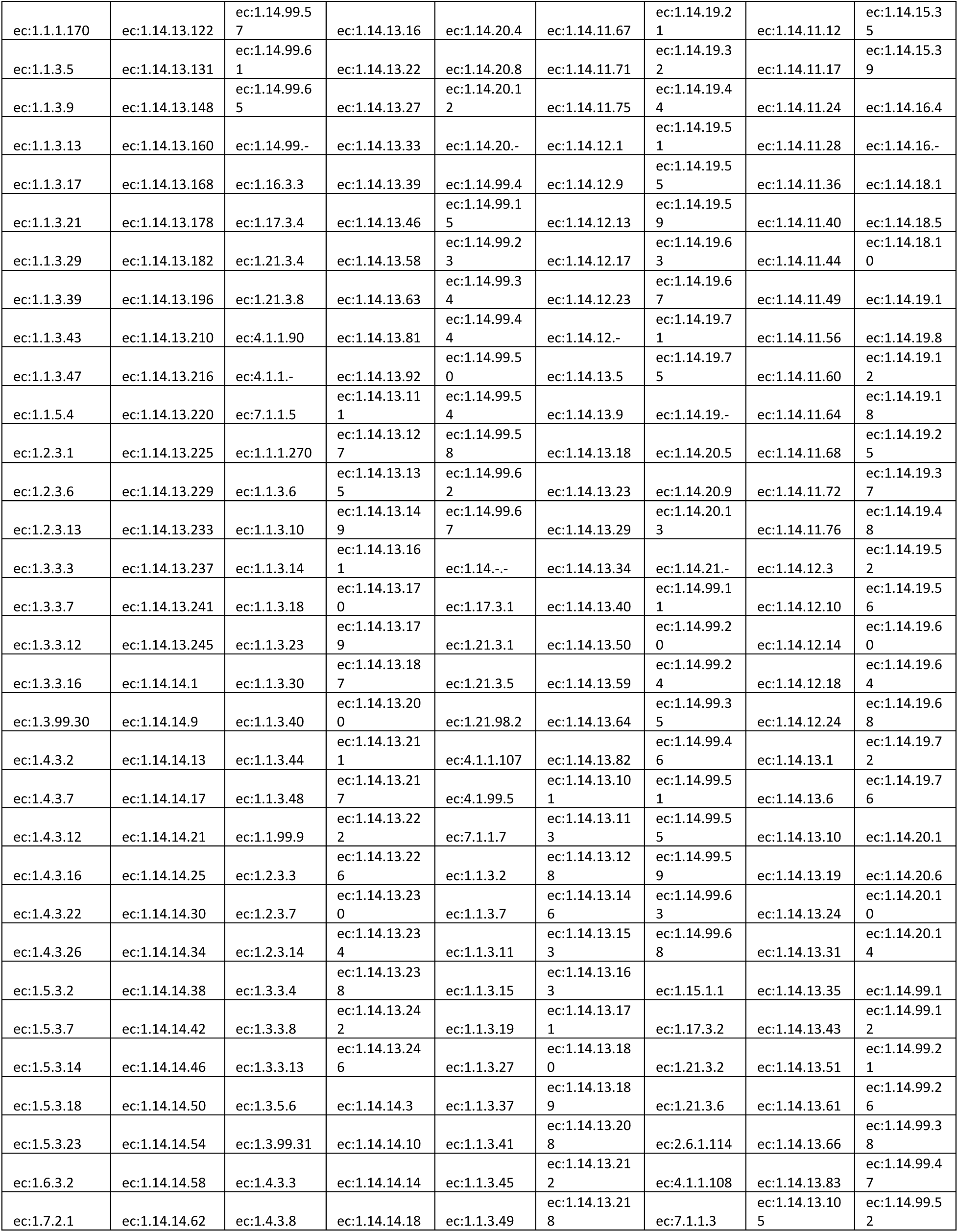
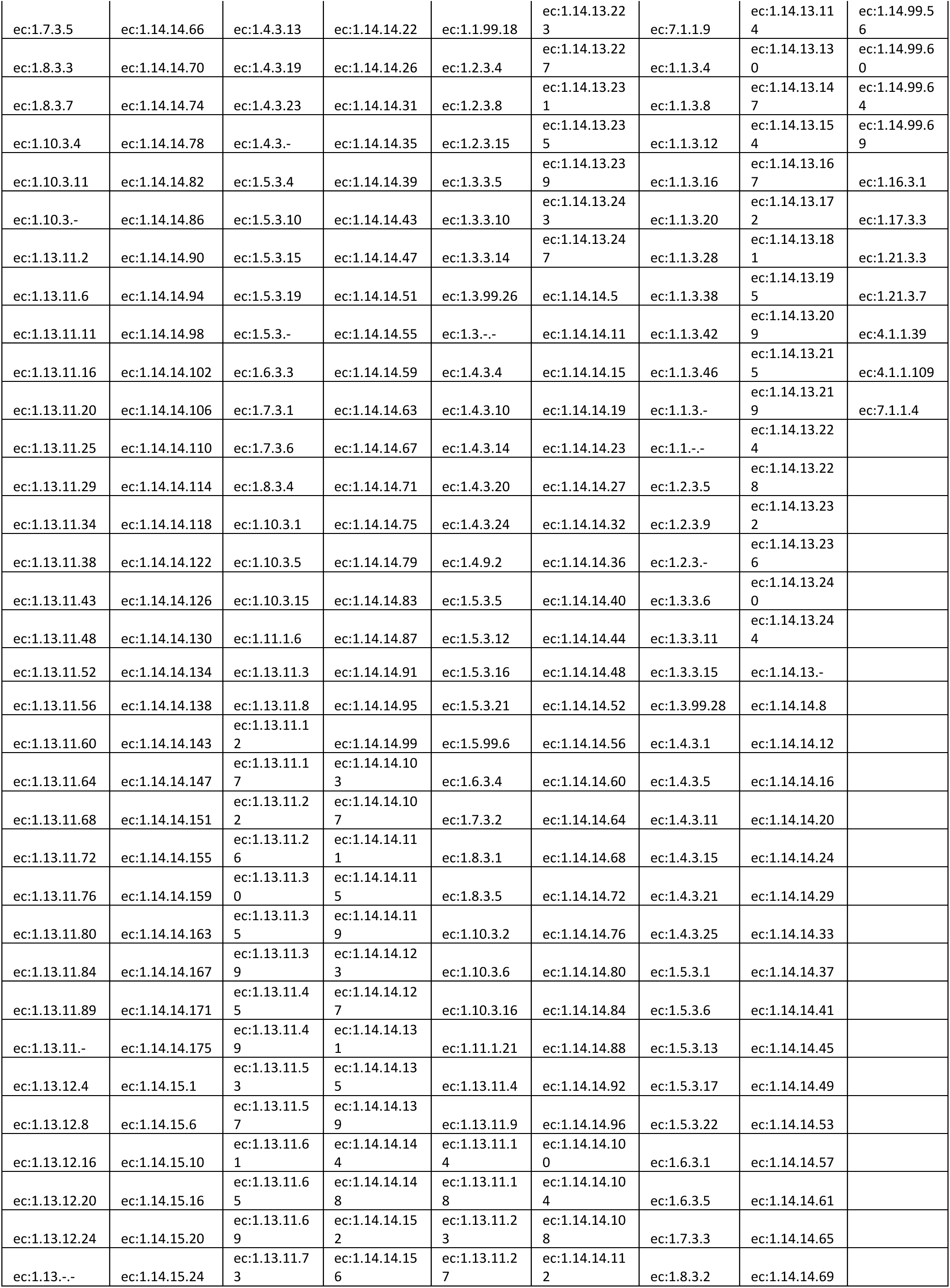
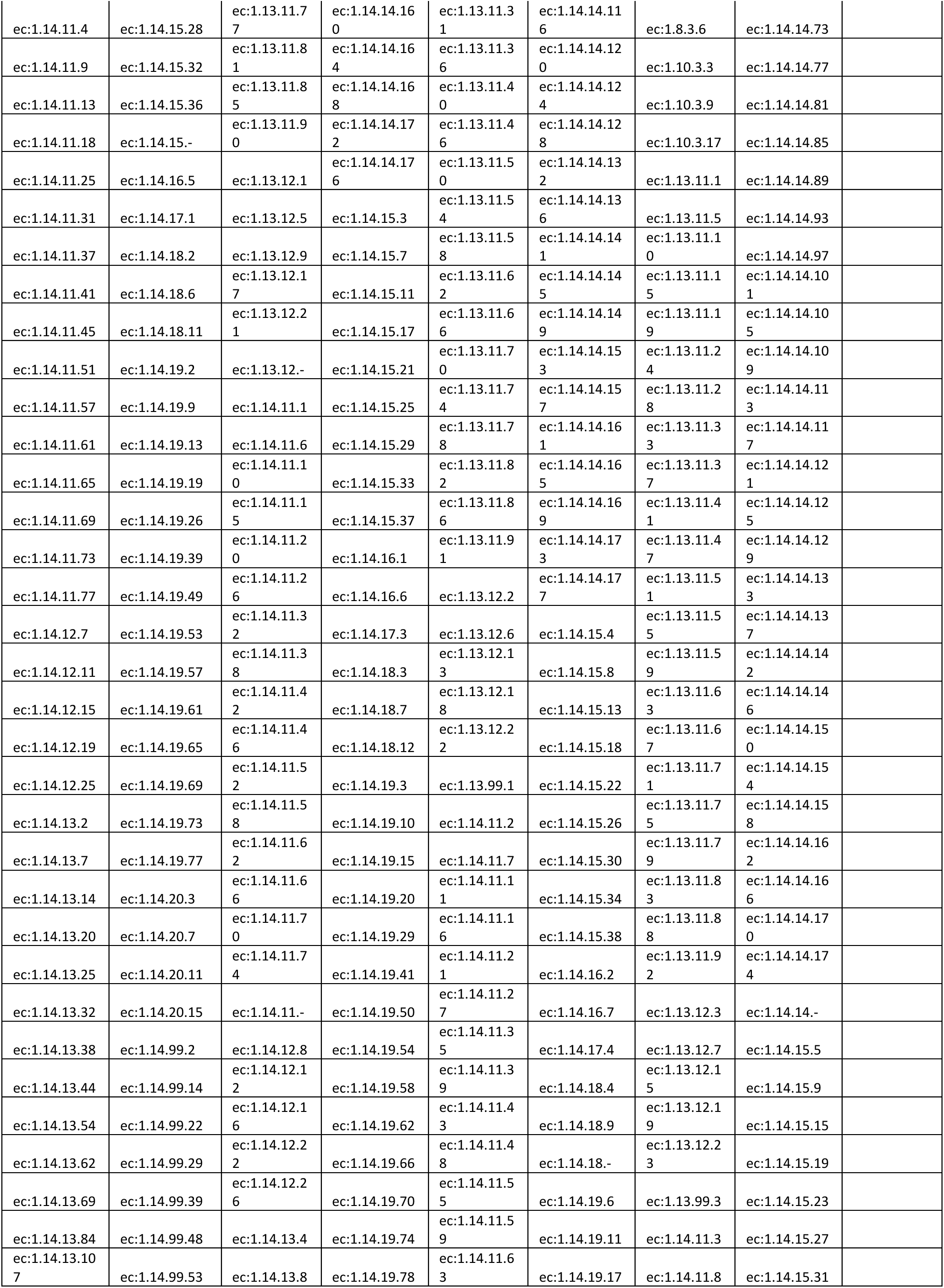
Enzymes that catalyze reactions which require molecular oxygen

**Table S4.** Master RClass similarity table with a cutoff value of 0.8

|  |  |  |  |
| --- | --- | --- | --- |
| RC00714 | 1 | RC00714 | O-Hydrolysis |
| RC00805 | 1 | RC00805 | De/hydrogenation |
| RC01114 | 1 | RC01114 | Anthocyanin/din di/hydroxylation |
| RC01553 | 1 | RC01553 | Anthocyanin/din de/hydrogenation |
| RC01632 | 1 | RC01632 | Di/hydroxylation |
| RC01837 | 1 | RC01837 | Di/hydroxylation |
| RC00046 | 1 | RC00046 | Di/hydroxylation |
| RC01587 | 1 | RC01587 | C-Hydrolysis |
| RC00234 | 1 | RC00234 | De/hydrogenation |
| RC00235 | 1 | RC00235 | De/hydrogenation |
| RC00657 | 1 | RC00657 | De/hydrogenation |
| RC00717 | 1 | RC00717 | De/hydrogenation |
| RC01235 | 1 | RC01235 | De/hydrogenation |
| RC00392 | 1 | RC00392 | De/methylation |
| RC00230 | 1 | RC00230 | Di/hydroxylation |
| RC00235 | 1 | RC00235 | Di/hydroxylation |
| RC00490 | 1 | RC00490 | Di/hydroxylation |
| RC01552 | 1 | RC01552 | Di/hydroxylation |
| RC00049 | 1 | RC00049 | O-Hydrolysis |
| RC00171 | 1 | RC00171 | O-Hydrolysis |
| RC00718 | 1 | RC00718 | Ring cleavage |
| RC02001 | 1 | RC02001 | Ring cleavage |
| RC02019 | 1 | RC02019 | Ring cleavage |
| RC02726 | 1 | RC02726 | Ring cleavage |
| <u>RC02046</u> | 0.9 | RC01632 | Di/hydroxylation |
| <u>RC00490</u> | 0.9 | RC00046 | Di/hydroxylation |
| <u>RC00236</u> | 0.9 | RC00046 | Di/hydroxylation |
| <u>RC00613</u> | 0.9 | RC00230 | Di/hydroxylation |
| <u>RC00046</u> | 0.9 | RC00490 | Di/hydroxylation |
| <u>RC02573</u> | 0.9 | RC00490 | Di/hydroxylation |
| <u>RC00589</u> | 0.9 | RC00490 | Di/hydroxylation |
| <u>RC01856</u> | 0.9 | RC00490 | Di/hydroxylation |
| <u>RC02016</u> | 0.9 | RC00490 | Di/hydroxylation |
| <u>RC00613</u> | 0.9 | RC01552 | Di/hydroxylation |
| <u>RC00274</u> | 0.9 | RC01552 | Di/hydroxylation |
| <u>RC00432</u> | 0.9 | RC01587 | C-Hydrolysis |
| <u>RC02019</u> | 0.9 | RC02001 | Ring cleavage |
| <u>RC02001</u> | 0.9 | RC02019 | Ring cleavage |
| <u>RC01053</u> | 0.877 | RC01632 | Di/hydroxylation |
| <u>RC01414</u> | 0.877 | RC01632 | Di/hydroxylation |
| <u>RC02573</u> | 0.867 | RC00046 | Di/hydroxylation |
| <u>RC00589</u> | 0.867 | RC00046 | Di/hydroxylation |
| <u>RC01856</u> | 0.867 | RC00046 | Di/hydroxylation |
| <u>RC02016</u> | 0.867 | RC00046 | Di/hydroxylation |
| <u>RC03365</u> | 0.867 | RC00046 | Di/hydroxylation |
| <u>RC01860</u> | 0.867 | RC00046 | Di/hydroxylation |
| <u>RC00661</u> | 0.867 | RC00230 | Di/hydroxylation |
| <u>RC00704</u> | 0.867 | RC00230 | Di/hydroxylation |
| <u>RC01993</u> | 0.867 | RC00230 | Di/hydroxylation |
| <u>RC00524</u> | 0.867 | RC00230 | Di/hydroxylation |
| <u>RC02085</u> | 0.867 | RC00230 | Di/hydroxylation |
| <u>RC01552</u> | 0.867 | RC00230 | Di/hydroxylation |
| <u>RC02734</u> | 0.867 | RC00230 | Di/hydroxylation |
| <u>RC03493</u> | 0.867 | RC00230 | Di/hydroxylation |
| <u>RC02043</u> | 0.867 | RC00230 | Di/hydroxylation |
| <u>RC00661</u> | 0.867 | RC01552 | Di/hydroxylation |
| <u>RC00277</u> | 0.867 | RC01552 | Di/hydroxylation |
| <u>RC01993</u> | 0.867 | RC01552 | Di/hydroxylation |
| <u>RC00524</u> | 0.867 | RC01552 | Di/hydroxylation |
| <u>RC02085</u> | 0.867 | RC01552 | Di/hydroxylation |
| <u>RC00230</u> | 0.867 | RC01552 | Di/hydroxylation |
| <u>RC02734</u> | 0.867 | RC01552 | Di/hydroxylation |
| <u>RC02043</u> | 0.867 | RC01552 | Di/hydroxylation |
| <u>RC03497</u> | 0.867 | RC01552 | Di/hydroxylation |
| <u>RC02955</u> | 0.867 | RC02726 | Ring cleavage |
| <u>RC02969</u> | 0.867 | RC02726 | Ring cleavage |
| <u>RC03232</u> | 0.86 | RC01632 | Di/hydroxylation |
| <u>RC00052</u> | 0.854 | RC01235 | De/hydrogenation |
| <u>RC02116</u> | 0.854 | RC01235 | De/hydrogenation |
| <u>RC00144</u> | 0.833 | RC00234 | De/hydrogenation |
| <u>RC00273</u> | 0.833 | RC00234 | De/hydrogenation |
| <u>RC00182</u> | 0.833 | RC00235 | De/hydrogenation |
| <u>RC00273</u> | 0.833 | RC00235 | De/hydrogenation |
| <u>RC02358</u> | 0.833 | RC01235 | De/hydrogenation |
| <u>RC01836</u> | 0.833 | RC00392 | De/methylation |
| <u>RC01916</u> | 0.833 | RC00392 | De/methylation |
| <u>RC01558</u> | 0.833 | RC00392 | De/methylation |
| <u>RC00505</u> | 0.833 | RC00046 | Di/hydroxylation |
| <u>RC00716</u> | 0.833 | RC00046 | Di/hydroxylation |
| <u>RC01316</u> | 0.833 | RC00046 | Di/hydroxylation |
| <u>RC00412</u> | 0.833 | RC00046 | Di/hydroxylation |
| <u>RC01801</u> | 0.833 | RC00046 | Di/hydroxylation |
| <u>RC00570</u> | 0.833 | RC00046 | Di/hydroxylation |
| <u>RC01919</u> | 0.833 | RC00046 | Di/hydroxylation |
| <u>RC00901</u> | 0.833 | RC00046 | Di/hydroxylation |
| <u>RC01836</u> | 0.833 | RC00046 | Di/hydroxylation |
| <u>RC00182</u> | 0.833 | RC00235 | Di/hydroxylation |
| <u>RC00273</u> | 0.833 | RC00235 | Di/hydroxylation |
| <u>RC01492</u> | 0.833 | RC01587 | C-Hydrolysis |
| <u>RC00452</u> | 0.833 | RC00049 | O-Hydrolysis |
| <u>RC01992</u> | 0.833 | RC00049 | O-Hydrolysis |
| <u>RC01669</u> | 0.833 | RC00049 | O-Hydrolysis |
| <u>RC01776</u> | 0.833 | RC00049 | O-Hydrolysis |
| <u>RC01684</u> | 0.833 | RC00171 | O-Hydrolysis |
| <u>RC02508</u> | 0.833 | RC00171 | O-Hydrolysis |
| <u>RC02023</u> | 0.833 | RC00171 | O-Hydrolysis |
| <u>RC02046</u> | 0.817 | RC01114 | Anthocyanin/din di/hydroxylation |
| <u>RC00466</u> | 0.8 | RC00714 | O-Hydrolysis |
| <u>RC00451</u> | 0.8 | RC00714 | O-Hydrolysis |
| <u>RC00746</u> | 0.8 | RC00714 | O-Hydrolysis |
| <u>RC01248</u> | 0.8 | RC00714 | O-Hydrolysis |
| <u>RC00049</u> | 0.8 | RC00714 | O-Hydrolysis |
| <u>RC00077</u> | 0.8 | RC00714 | O-Hydrolysis |
| <u>RC00350</u> | 0.8 | RC00714 | O-Hydrolysis |
| <u>RC00233</u> | 0.8 | RC00714 | O-Hydrolysis |
| <u>RC00878</u> | 0.8 | RC00714 | O-Hydrolysis |
| <u>RC00530</u> | 0.8 | RC00714 | O-Hydrolysis |
| <u>RC00231</u> | 0.8 | RC00714 | O-Hydrolysis |
| <u>RC02123</u> | 0.8 | RC00392 | De/methylation |
| <u>RC01746</u> | 0.8 | RC00392 | De/methylation |
| <u>RC02082</u> | 0.8 | RC00392 | De/methylation |
| <u>RC00466</u> | 0.8 | RC00392 | De/methylation |
| <u>RC01324</u> | 0.8 | RC00392 | De/methylation |
| <u>RC01895</u> | 0.8 | RC00392 | De/methylation |
| <u>RC01372</u> | 0.8 | RC00392 | De/methylation |
| <u>RC03192</u> | 0.8 | RC00392 | De/methylation |
| <u>RC00171</u> | 0.8 | RC00392 | De/methylation |
| <u>RC00878</u> | 0.8 | RC00392 | De/methylation |
| <u>RC00128</u> | 0.8 | RC00392 | De/methylation |
| <u>RC00332</u> | 0.8 | RC00046 | Di/hydroxylation |
| <u>RC00209</u> | 0.8 | RC00046 | Di/hydroxylation |
| <u>RC00701</u> | 0.8 | RC00046 | Di/hydroxylation |
| <u>RC03141</u> | 0.8 | RC00046 | Di/hydroxylation |
| <u>RC00432</u> | 0.8 | RC00046 | Di/hydroxylation |
| <u>RC01590</u> | 0.8 | RC00046 | Di/hydroxylation |
| <u>RC01921</u> | 0.8 | RC00046 | Di/hydroxylation |
| <u>RC00390</u> | 0.8 | RC00046 | Di/hydroxylation |
| <u>RC01838</u> | 0.8 | RC00046 | Di/hydroxylation |
| <u>RC02112</u> | 0.8 | RC00046 | Di/hydroxylation |
| <u>RC01199</u> | 0.8 | RC00046 | Di/hydroxylation |
| <u>RC03483</u> | 0.8 | RC00046 | Di/hydroxylation |
| <u>RC01913</u> | 0.8 | RC00046 | Di/hydroxylation |
| <u>RC00949</u> | 0.8 | RC00046 | Di/hydroxylation |
| <u>RC03265</u> | 0.8 | RC00230 | Di/hydroxylation |
| <u>RC00478</u> | 0.8 | RC00230 | Di/hydroxylation |
| <u>RC01218</u> | 0.8 | RC00230 | Di/hydroxylation |
| <u>RC01963</u> | 0.8 | RC00230 | Di/hydroxylation |
| <u>RC00723</u> | 0.8 | RC00230 | Di/hydroxylation |
| <u>RC00277</u> | 0.8 | RC00230 | Di/hydroxylation |
| <u>RC02241</u> | 0.8 | RC00230 | Di/hydroxylation |
| <u>RC01203</u> | 0.8 | RC00230 | Di/hydroxylation |
| <u>RC01480</u> | 0.8 | RC00230 | Di/hydroxylation |
| <u>RC02731</u> | 0.8 | RC00230 | Di/hydroxylation |
| <u>RC01592</u> | 0.8 | RC00230 | Di/hydroxylation |
| <u>RC00274</u> | 0.8 | RC00230 | Di/hydroxylation |
| <u>RC03497</u> | 0.8 | RC00230 | Di/hydroxylation |
| <u>RC00364</u> | 0.8 | RC00490 | Di/hydroxylation |
| <u>RC00391</u> | 0.8 | RC00490 | Di/hydroxylation |
| <u>RC00236</u> | 0.8 | RC00490 | Di/hydroxylation |
| <u>RC03365</u> | 0.8 | RC00490 | Di/hydroxylation |
| <u>RC01860</u> | 0.8 | RC00490 | Di/hydroxylation |
| <u>RC01637</u> | 0.8 | RC00490 | Di/hydroxylation |
| <u>RC00478</u> | 0.8 | RC01552 | Di/hydroxylation |
| <u>RC01218</u> | 0.8 | RC01552 | Di/hydroxylation |
| <u>RC01963</u> | 0.8 | RC01552 | Di/hydroxylation |
| <u>RC00723</u> | 0.8 | RC01552 | Di/hydroxylation |
| <u>RC00704</u> | 0.8 | RC01552 | Di/hydroxylation |
| <u>RC02241</u> | 0.8 | RC01552 | Di/hydroxylation |
| <u>RC01203</u> | 0.8 | RC01552 | Di/hydroxylation |
| <u>RC01480</u> | 0.8 | RC01552 | Di/hydroxylation |
| <u>RC03493</u> | 0.8 | RC01552 | Di/hydroxylation |
| <u>RC02731</u> | 0.8 | RC01552 | Di/hydroxylation |
| <u>RC01592</u> | 0.8 | RC01552 | Di/hydroxylation |
| <u>RC01662</u> | 0.8 | RC01587 | C-Hydrolysis |
| <u>RC01869</u> | 0.8 | RC01587 | C-Hydrolysis |
| <u>RC03346</u> | 0.8 | RC01587 | C-Hydrolysis |
| <u>RC02379</u> | 0.8 | RC01587 | C-Hydrolysis |
| <u>RC00801</u> | 0.8 | RC01587 | C-Hydrolysis |
| <u>RC00569</u> | 0.8 | RC01587 | C-Hydrolysis |
| <u>RC00236</u> | 0.8 | RC01587 | C-Hydrolysis |
| <u>RC01363</u> | 0.8 | RC01587 | C-Hydrolysis |
| <u>RC00466</u> | 0.8 | RC00049 | O-Hydrolysis |
| <u>RC00451</u> | 0.8 | RC00049 | O-Hydrolysis |
| <u>RC00746</u> | 0.8 | RC00049 | O-Hydrolysis |
| <u>RC01248</u> | 0.8 | RC00049 | O-Hydrolysis |
| <u>RC00059</u> | 0.8 | RC00049 | O-Hydrolysis |
| <u>RC00397</u> | 0.8 | RC00049 | O-Hydrolysis |
| <u>RC00262</u> | 0.8 | RC00049 | O-Hydrolysis |
| <u>RC00028</u> | 0.8 | RC00049 | O-Hydrolysis |
| <u>RC02398</u> | 0.8 | RC00049 | O-Hydrolysis |
| <u>RC00171</u> | 0.8 | RC00049 | O-Hydrolysis |
| <u>RC00529</u> | 0.8 | RC00049 | O-Hydrolysis |
| <u>RC00077</u> | 0.8 | RC00049 | O-Hydrolysis |
| <u>RC00350</u> | 0.8 | RC00049 | O-Hydrolysis |
| <u>RC00714</u> | 0.8 | RC00049 | O-Hydrolysis |
| <u>RC00530</u> | 0.8 | RC00049 | O-Hydrolysis |
| <u>RC00231</u> | 0.8 | RC00049 | O-Hydrolysis |
| <u>RC00392</u> | 0.8 | RC00171 | O-Hydrolysis |
| <u>RC01372</u> | 0.8 | RC00171 | O-Hydrolysis |
| <u>RC03192</u> | 0.8 | RC00171 | O-Hydrolysis |
| <u>RC00059</u> | 0.8 | RC00171 | O-Hydrolysis |
| <u>RC00397</u> | 0.8 | RC00171 | O-Hydrolysis |
| <u>RC00262</u> | 0.8 | RC00171 | O-Hydrolysis |
| <u>RC00049</u> | 0.8 | RC00171 | O-Hydrolysis |
| <u>RC00028</u> | 0.8 | RC00171 | O-Hydrolysis |
| <u>RC02398</u> | 0.8 | RC00171 | O-Hydrolysis |
| <u>RC00529</u> | 0.8 | RC00171 | O-Hydrolysis |
| <u>RC00878</u> | 0.8 | RC00171 | O-Hydrolysis |
| <u>RC00128</u> | 0.8 | RC00171 | O-Hydrolysis |

## Source Codes

### Input: Enzyme

Below code can take an enzyme in the form of EC number and outputs a list of predictions. Example input: enzyme = ‘ec:2.3.1.74’

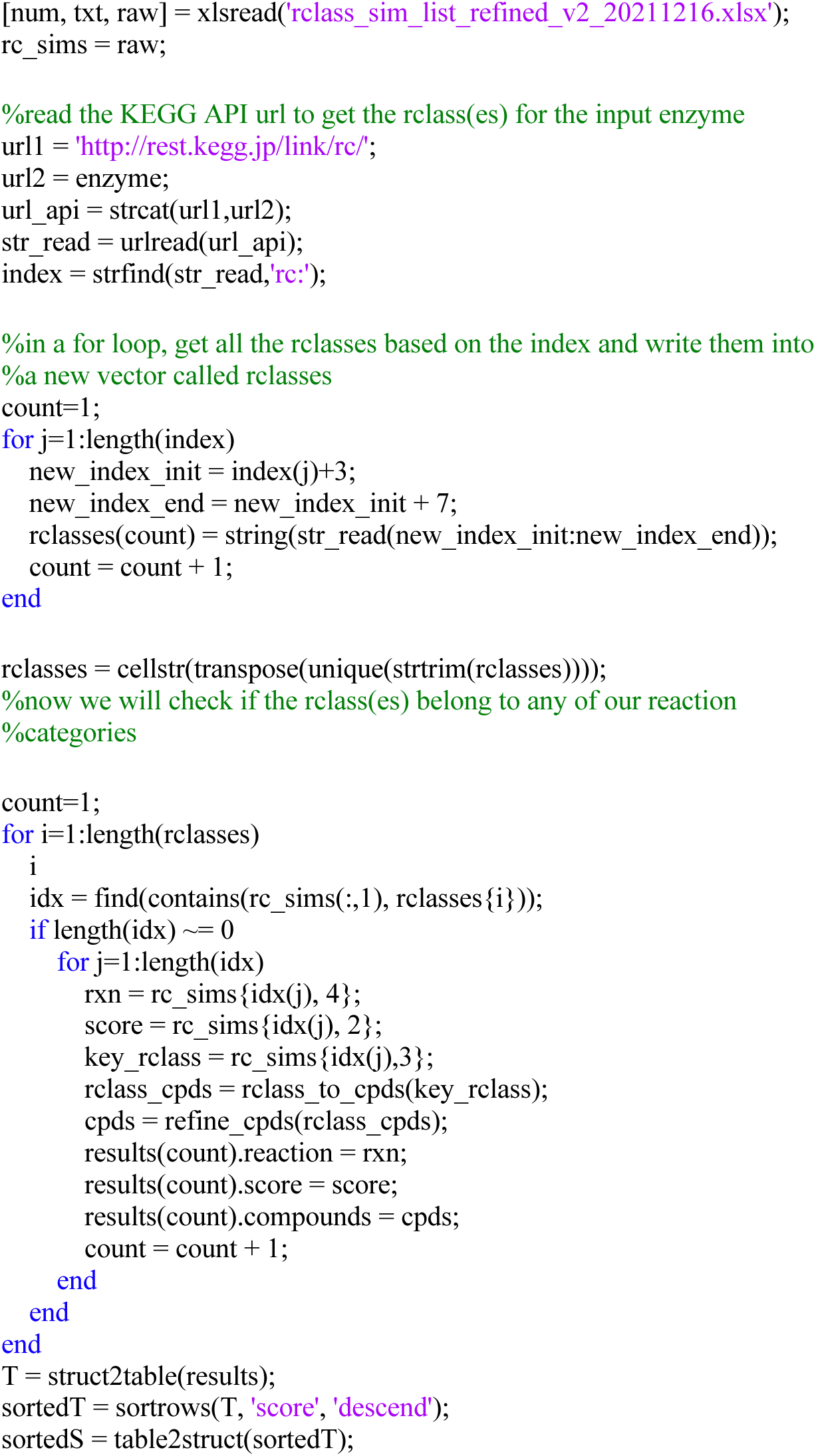

### Input: Organism

Below code can take an organism in the form of KEGG organism ID and outputs a list of predictions. Example input: mic = ‘fpla’

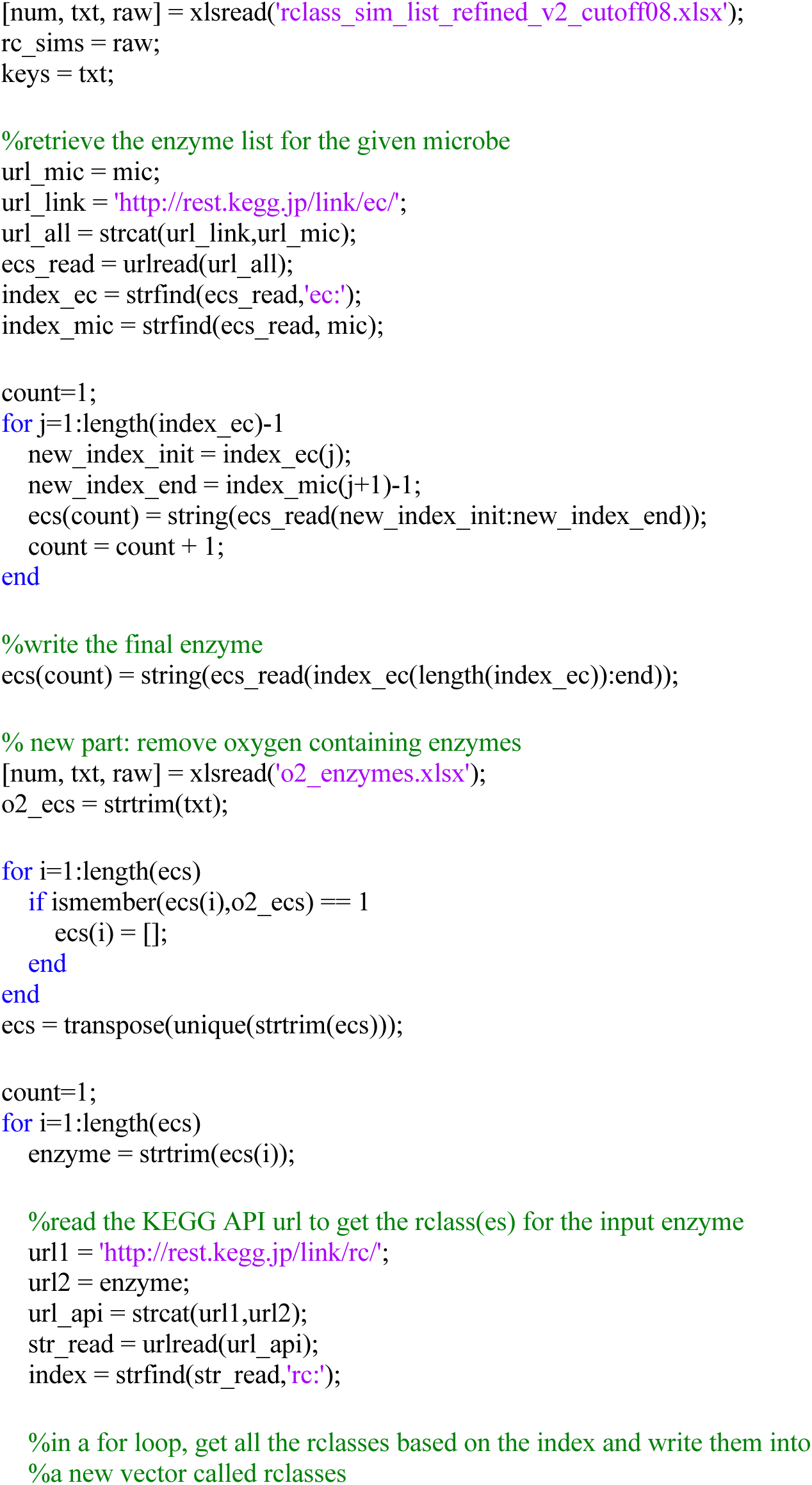
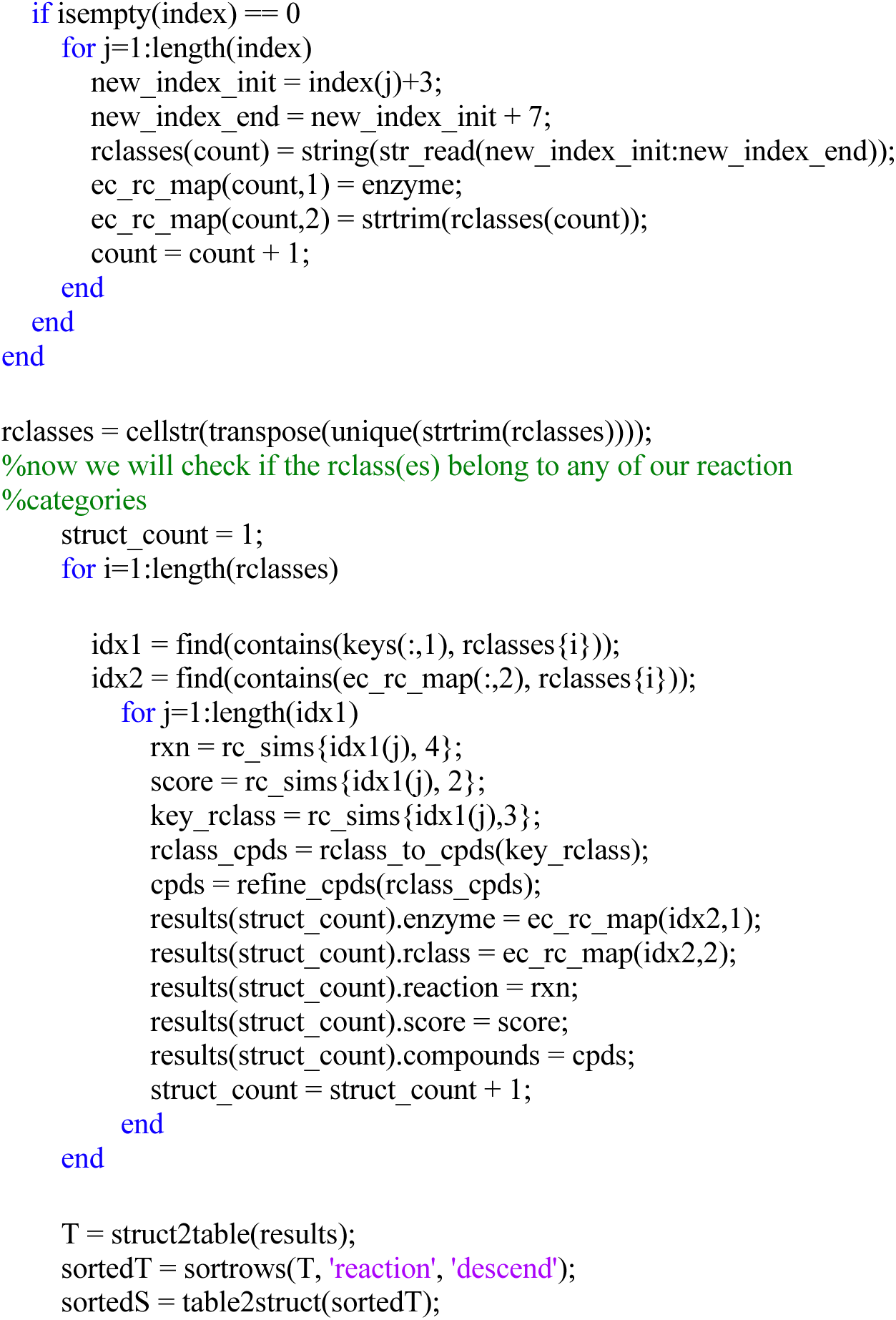

## Subfunctions

### Retrieving the compounds from a given RClass

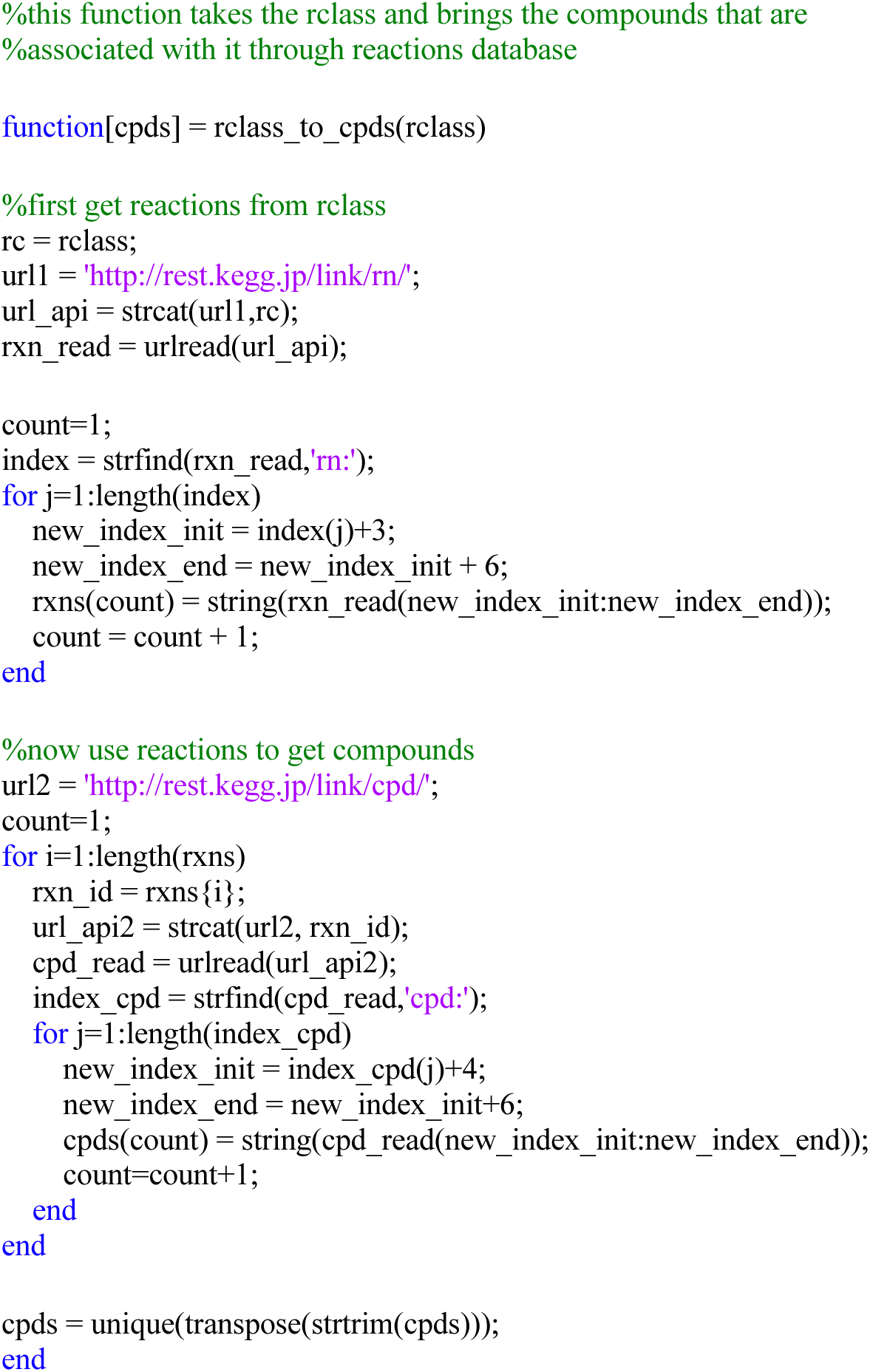

### Refining the compounds to obtain only flavonoids and their first derivatives

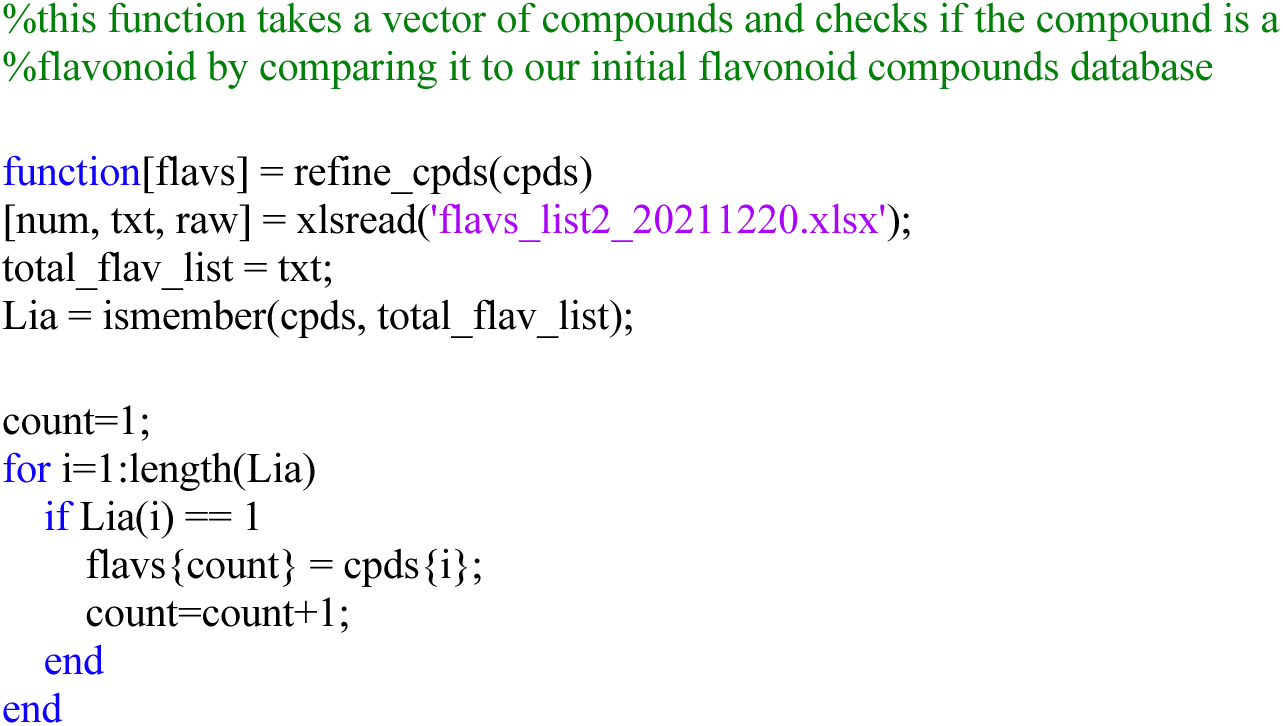

## Notes

### Competing Interest Statement

The authors have declared no competing interest.

## References

1. Amin, S. A., Chavez, E., Porokhin, V., Nair, N. U., & Hassoun, S. (2019). Towards creating an extended metabolic model (EMM) for E. coli using enzyme promiscuity prediction and metabolomics data. Microb Cell Fact, 18(1), 109. https://doi.org/10.1186/s12934-019-1156-3

2. Anhe, F. F., Varin, T. V., Le Barz, M., Desjardins, Y., Levy, E., Roy, D., & Marette, A. (2015). Gut Microbiota Dysbiosis in Obesity-Linked Metabolic Diseases and Prebiotic Potential of Polyphenol-Rich Extracts. Curr Obes Rep, 4(4), 389–400. https://doi.org/10.1007/s13679-015-0172-9

3. Ávila, M., Hidalgo, M., Sánchez-Moreno, C., Pelaez, C., Requena, T., & de Pascual-Teresa, S. (2009). Bioconversion of anthocyanin glycosides by Bifidobacteria and Lactobacillus. Food research international, 42(10), 1453–1461.

4. Axelson, M., & Setchell, K. D. (1981). The excretion of lignans in rats -- evidence for an intestinal bacterial source for this new group of compounds. FEBS Lett, 123(2), 337–342. https://doi.org/10.1016/0014-5793(81)80322-5

5. Baas, B. J., Zandvoort, E., Geertsema, E. M., & Poelarends, G. J. (2013). Recent advances in the study of enzyme promiscuity in the tautomerase superfamily. Chembiochem, 14(8), 917–926. https://doi.org/10.1002/cbic.201300098

6. Braune, A., Engst, W., Elsinghorst, P. W., Furtmann, N., Bajorath, J., Gutschow, M., & Blaut, M. (2016). Chalcone Isomerase from Eubacterium ramulus Catalyzes the Ring Contraction of Flavanonols. J Bacteriol, 198(21), 2965–2974. https://doi.org/10.1128/JB.00490-16

7. Braune, A., Gutschow, M., & Blaut, M. (2019). An NADH-Dependent Reductase from Eubacterium ramulus Catalyzes the Stereospecific Heteroring Cleavage of Flavanones and Flavanonols. Appl Environ Microbiol, 85(19). https://doi.org/10.1128/AEM.01233-19

8. Bravo, J. A., Forsythe, P., Chew, M. V., Escaravage, E., Savignac, H. M., Dinan, T. G., Bienenstock, J., & Cryan, J. F. (2011). Ingestion of Lactobacillus strain regulates emotional behavior and central GABA receptor expression in a mouse via the vagus nerve. Proc Natl Acad Sci U S A, 108(38), 16050–16055. https://doi.org/10.1073/pnas.1102999108

9. Carbonell, P., Wong, J., Swainston, N., Takano, E., Turner, N. J., Scrutton, N. S., Kell, D. B., Breitling, R., & Faulon, J. L. (2018). Selenzyme: enzyme selection tool for pathway design. Bioinformatics, 34(12), 2153–2154. https://doi.org/10.1093/bioinformatics/bty065

10. Carrera-Quintanar, L., Lopez Roa, R. I., Quintero-Fabian, S., Sanchez-Sanchez, M. A., Vizmanos, B., & Ortuno-Sahagun, D. (2018). Phytochemicals That Influence Gut Microbiota as Prophylactics and for the Treatment of Obesity and Inflammatory Diseases. Mediators Inflamm, 2018, 9734845. https://doi.org/10.1155/2018/9734845

11. Caspi, R., Billington, R., Keseler, I. M., Kothari, A., Krummenacker, M., Midford, P. E., Ong, W. K., Paley, S., Subhraveti, P., & Karp, P. D. (2020). The MetaCyc database of metabolic pathways and enzymes - a 2019 update. Nucleic Acids Res, 48(D1), D445–D453. https://doi.org/10.1093/nar/gkz862

12. Cattaneo, A., Cattane, N., Galluzzi, S., Provasi, S., Lopizzo, N., Festari, C., Ferrari, C., Guerra, U. P., Paghera, B., Muscio, C., Bianchetti, A., Volta, G. D., Turla, M., Cotelli, M. S., Gennuso, M., Prelle, A., Zanetti, O., Lussignoli, G., Mirabile, D., . . . Group, I.-F. (2017). Association of brain amyloidosis with pro-inflammatory gut bacterial taxa and peripheral inflammation markers in cognitively impaired elderly. Neurobiol Aging, 49, 60–68. https://doi.org/10.1016/j.neurobiolaging.2016.08.019

13. Chen, J., Yang, J., Ma, L., Li, J., Shahzad, N., & Kim, C. K. (2020). Structure-antioxidant activity relationship of methoxy, phenolic hydroxyl, and carboxylic acid groups of phenolic acids. Sci Rep, 10(1), 2611. https://doi.org/10.1038/s41598-020-59451-z

14. Darsandhari, S., Dhakal, D., Shrestha, B., Parajuli, P., Seo, J. H., Kim, T. S., & Sohng, J. K. (2018). Characterization of regioselective flavonoid O-methyltransferase from the Streptomyces sp. KCTC 0041BP. Enzyme Microb Technol, 113, 29–36. https://doi.org/10.1016/j.enzmictec.2018.02.007

15. Delepine, B., Duigou, T., Carbonell, P., & Faulon, J. L. (2018). RetroPath2.0: A retrosynthesis workflow for metabolic engineers. Metab Eng, 45, 158–170. https://doi.org/10.1016/j.ymben.2017.12.002

16. Finnigan, W., Hepworth, L. J., Flitsch, S. L., & Turner, N. J. (2021). RetroBioCat as a computer-aided synthesis planning tool for biocatalytic reactions and cascades. Nat Catal, 4(2), 98–104. https://doi.org/10.1038/s41929-020-00556-z

17. Frankenfeld, C. L. (2011). O-desmethylangolensin: the importance of equol’s lesser known cousin to human health. Adv Nutr, 2(4), 317–324. https://doi.org/10.3945/an.111.000539

18. Galvez, J., Estrada-Reyes, R., Benitez-King, G., Araujo, G., Orozco, S., Fernandez-Mas, R., Almazan, S., & Calixto, E. (2015). Involvement of the GABAergic system in the neuroprotective and sedative effects of acacetin 7-O-glucoside in rodents. Restor Neurol Neurosci, 33(5), 683–700. https://doi.org/10.3233/RNN-140486

19. Gardana, C., Canzi, E., & Simonetti, P. (2014). R(-)-O-desmethylangolensin is the main enantiomeric form of daidzein metabolite produced by human in vitro and in vivo. J Chromatogr B Analyt Technol Biomed Life Sci, 953–954, 30-37. https://doi.org/10.1016/j.jchromb.2014.01.048

20. Gulsan, E. E., Nowshad, F., Jayaraman, A., & Lee, K. (2022). Metabolism of Dietary Carbohydrates by Intestinal Bacteria. In Metabolism of Nutrients by Gut Microbiota (pp. 18–47).

21. Gülşan, E. E., Nowshad, F., Leigh, M. D., Crott, J. W., Safe, S., Jayaraman, A., & Lee, K. (2022). A Chalcone Synthase-Like Bacterial Protein Catalyzes Heterocyclic C-Ring Cleavage of Naringenin to Alter Bioactivity Against Nuclear Receptors in Colonic Epithelial Cells. bioRxiv, 2022.2004. 2022.489210.

22. Hafner, J., Payne, J., MohammadiPeyhani, H., Hatzimanikatis, V., & Smolke, C. (2021). A computational workflow for the expansion of heterologous biosynthetic pathways to natural product derivatives. Nat Commun, 12(1), 1760. https://doi.org/10.1038/s41467-021-22022-5

23. Houlden, A., Goldrick, M., Brough, D., Vizi, E. S., Lenart, N., Martinecz, B., Roberts, I. S., & Denes, A. (2016). Brain injury induces specific changes in the caecal microbiota of mice via altered autonomic activity and mucoprotein production. Brain Behav Immun, 57, 10–20. https://doi.org/10.1016/j.bbi.2016.04.003

24. Jackson, R. L., Greiwe, J. S., & Schwen, R. J. (2011). Emerging evidence of the health benefits of S-equol, an estrogen receptor beta agonist. Nutr Rev, 69(8), 432–448. https://doi.org/10.1111/j.1753-4887.2011.00400.x

25. Jeffryes, J. G., Seaver, S. M. D., Faria, J. P., & Henry, C. S. (2018). A pathway for every product? Tools to discover and design plant metabolism. Plant Sci, 273, 61–70. https://doi.org/10.1016/j.plantsci.2018.03.025

26. Jiang, H., Fang, J., Xing, J., Wang, L., Wang, Q., Wang, Y., Li, Z., & Liu, R. (2019). Tilianin mediates neuroprotection against ischemic injury by attenuating CaMKII-dependent mitochondrion-mediated apoptosis and MAPK/NF-kappaB signaling. Life Sci, 216, 233–245. https://doi.org/10.1016/j.lfs.2018.11.035

27. Jiao, X., Wang, Y., Lin, Y., Lang, Y., Li, E., Zhang, X., Zhang, Q., Feng, Y., Meng, X., & Li, B. (2019). Blueberry polyphenols extract as a potential prebiotic with anti-obesity effects on C57BL/6 J mice by modulating the gut microbiota. J Nutr Biochem, 64, 88–100. https://doi.org/10.1016/j.jnutbio.2018.07.008

28. Judson, P. N., Long, A., Murray, E., & Patel, M. (2015). Assessing Confidence in Predictions Using Veracity and Utility - A Case Study on the Prediction of Mammalian Metabolism by Meteor Nexus. Mol Inform, 34(5), 284–291. https://doi.org/10.1002/minf.201400184

29. Kanehisa, M., & Goto, S. (2000). KEGG: kyoto encyclopedia of genes and genomes. Nucleic Acids Res, 28(1), 27–30. https://doi.org/10.1093/nar/28.1.27

30. Khattulanuar, F. S., Sekar, M., Fuloria, S., Gan, S. H., Rani, N., Ravi, S., Chidambaram, K., Begum, M. Y., Azad, A. K., Jeyabalan, S., Dhiravidamani, A., Thangavelu, L., Lum, P. T., Subramaniyan, V., Wu, Y. S., Sathasivam, K. V., & Fuloria, N. K. (2022). Tilianin: A Potential Natural Lead Molecule for New Drug Design and Development for the Treatment of Cardiovascular Disorders. Molecules, 27(3). https://doi.org/10.3390/molecules27030673

31. Khersonsky, O., & Tawfik, D. S. (2010). Enzyme promiscuity: a mechanistic and evolutionary perspective. Annu Rev Biochem, 79, 471–505. https://doi.org/10.1146/annurev-biochem-030409-143718

32. Kim, M., Kim, N., & Han, J. (2014). Metabolism of Kaempferia parviflora polymethoxyflavones by human intestinal bacterium Bautia sp. MRG-PMF1. J Agric Food Chem, 62(51), 12377–12383. https://doi.org/10.1021/jf504074n

33. Kim, M. S., Kim, Y., Choi, H., Kim, W., Park, S., Lee, D., Kim, D. K., Kim, H. J., Choi, H., Hyun, D. W., Lee, J. Y., Choi, E. Y., Lee, D. S., Bae, J. W., & Mook-Jung, I. (2020). Transfer of a healthy microbiota reduces amyloid and tau pathology in an Alzheimer’s disease animal model. Gut, 69(2), 283–294. https://doi.org/10.1136/gutjnl-2018-317431

34. Koch, M., Duigou, T., & Faulon, J. L. (2020). Reinforcement Learning for Bioretrosynthesis. ACS Synth Biol, 9(1), 157–168. https://doi.org/10.1021/acssynbio.9b00447

35. Kuwahara, H., Alazmi, M., Cui, X., & Gao, X. (2016). MRE: a web tool to suggest foreign enzymes for the biosynthesis pathway design with competing endogenous reactions in mind. Nucleic Acids Res, 44(W1), W217–225. https://doi.org/10.1093/nar/gkw342

36. Lagkouvardos, I., Pukall, R., Abt, B., Foesel, B. U., Meier-Kolthoff, J. P., Kumar, N., Bresciani, A., Martinez, I., Just, S., Ziegler, C., Brugiroux, S., Garzetti, D., Wenning, M., Bui, T. P., Wang, J., Hugenholtz, F., Plugge, C. M., Peterson, D. A., Hornef, M. W., . . . Clavel, T. (2016). The Mouse Intestinal Bacterial Collection (miBC) provides host-specific insight into cultured diversity and functional potential of the gut microbiota. Nat Microbiol, 1(10), 16131. https://doi.org/10.1038/nmicrobiol.2016.131

37. Latendresse, M., Krummenacker, M., & Karp, P. D. (2014). Optimal metabolic route search based on atom mappings. Bioinformatics, 30(14), 2043–2050. https://doi.org/10.1093/bioinformatics/btu150

38. Leinonen, R., Sugawara, H., Shumway, M., & International Nucleotide Sequence Database, C. (2011). The sequence read archive. Nucleic Acids Res, 39(Database issue), D19–21. https://doi.org/10.1093/nar/gkq1019

39. Li, C., Lee, M. J., Sheng, S., Meng, X., Prabhu, S., Winnik, B., Huang, B., Chung, J. Y., Yan, S., Ho, C. T., & Yang, C. S. (2000). Structural identification of two metabolites of catechins and their kinetics in human urine and blood after tea ingestion. Chem Res Toxicol, 13(3), 177–184. https://doi.org/10.1021/tx9901837

40. Liu, B., Ramsundar, B., Kawthekar, P., Shi, J., Gomes, J., Luu Nguyen, Q., Ho, S., Sloane, J., Wender, P., & Pande, V. (2017). Retrosynthetic Reaction Prediction Using Neural Sequence-to-Sequence Models. ACS Cent Sci, 3(10), 1103–1113. https://doi.org/10.1021/acscentsci.7b00303

41. Lund, T. D., Munson, D. J., Haldy, M. E., Setchell, K. D., Lephart, E. D., & Handa, R. J. (2004). Equol is a novel anti-androgen that inhibits prostate growth and hormone feedback. Biol Reprod, 70(4), 1188–1195. https://doi.org/10.1095/biolreprod.103.023713

42. Marotti, I., Bonetti, A., Biavati, B., Catizone, P., & Dinelli, G. (2007). Biotransformation of common bean (Phaseolus vulgaris L.) flavonoid glycosides by bifidobacterium species from human intestinal origin. J Agric Food Chem, 55(10), 3913–3919. https://doi.org/10.1021/jf062997g

43. Mayo, B., Vazquez, L., & Florez, A. B. (2019). Equol: A Bacterial Metabolite from The Daidzein Isoflavone and Its Presumed Beneficial Health Effects. Nutrients, 11(9). https://doi.org/10.3390/nu11092231

44. McGuigan, J. E., Chang, Y., & Dajani, E. Z. (1986). Effect of misoprostol, an antiulcer prostaglandin, on serum gastrin in patients with duodenal ulcer. Digestive Diseases and Sciences, 31, 120S–125S.

45. Morishita, S., Onomura, T., Inoue, T., Maeda, H., & Akagi, H. (1989). Bone scintigraphy in patients with breast cancer, pulmonary cancer, uterine cervix cancer, and prostatic cancer. Statistical study of spinal accumulation cases. Spine (Phila Pa 1976), 14(8), 784–789. https://doi.org/10.1097/00007632-198908000-00002

46. Muto, A., Kotera, M., Tokimatsu, T., Nakagawa, Z., Goto, S., & Kanehisa, M. (2013). Modular architecture of metabolic pathways revealed by conserved sequences of reactions. J Chem Inf Model, 53(3), 613–622. https://doi.org/10.1021/ci3005379

47. Nam, K. H., Choi, J. H., Seo, Y. J., Lee, Y. M., Won, Y. S., Lee, M. R., Lee, M. N., Park, J. G., Kim, Y. M., Kim, H. C., Lee, C. H., Lee, H. K., Oh, S. R., & Oh, G. T. (2006). Inhibitory effects of tilianin on the expression of inducible nitric oxide synthase in low density lipoprotein receptor deficiency mice. Exp Mol Med, 38(4), 445–452. https://doi.org/10.1038/emm.2006.52

48. Panche, A. N., Diwan, A. D., & Chandra, S. R. (2016). Flavonoids: an overview. J Nutr Sci, 5, e47. https://doi.org/10.1017/jns.2016.41

49. Ponzetto, A., Negro, F., Popper, H., Bonino, F., Engle, R., Rizzetto, M., Purcell, R. H., & Gerin, J. L. (1988). Serial passage of hepatitis delta virus in chronic hepatitis B virus carrier chimpanzees. Hepatology, 8(6), 1655–1661. https://doi.org/10.1002/hep.1840080631

50. Saad, M. J., Santos, A., & Prada, P. O. (2016). Linking Gut Microbiota and Inflammation to Obesity and Insulin Resistance. Physiology (Bethesda), 31(4), 283–293. https://doi.org/10.1152/physiol.00041.2015

51. Schmidt, A., Li, C., Jones, A. D., & Pichersky, E. (2012). Characterization of a flavonol 3-O-methyltransferase in the trichomes of the wild tomato species Solanum habrochaites. Planta, 236(3), 839–849. https://doi.org/10.1007/s00425-012-1676-0

52. Schneider, H., & Blaut, M. (2000). Anaerobic degradation of flavonoids by Eubacterium ramulus. Arch Microbiol, 173(1), 71–75. https://doi.org/10.1007/s002030050010

53. Schoefer, L., Braune, A., & Blaut, M. (2004). Cloning and expression of a phloretin hydrolase gene from Eubacterium ramulus and characterization of the recombinant enzyme. Appl Environ Microbiol, 70(10), 6131–6137. https://doi.org/10.1128/AEM.70.10.6131-6137.2004

54. Schomburg, I., Chang, A., Ebeling, C., Gremse, M., Heldt, C., Huhn, G., & Schomburg, D. (2004). BRENDA, the enzyme database: updates and major new developments. Nucleic Acids Res, 32(Database issue), D431–433. https://doi.org/10.1093/nar/gkh081

55. Setchell, K. D., Clerici, C., Lephart, E. D., Cole, S. J., Heenan, C., Castellani, D., Wolfe, B. E., Nechemias-Zimmer, L., Brown, N. M., Lund, T. D., Handa, R. J., & Heubi, J. E. (2005). S-equol, a potent ligand for estrogen receptor beta, is the exclusive enantiomeric form of the soy isoflavone metabolite produced by human intestinal bacterial flora. Am J Clin Nutr, 81(5), 1072–1079. https://doi.org/10.1093/ajcn/81.5.1072

56. Singh, A., Kukreti, R., Saso, L., & Kukreti, S. (2019). Oxidative Stress: A Key Modulator in Neurodegenerative Diseases. Molecules, 24(8). https://doi.org/10.3390/molecules24081583

57. UniProt, C. (2021). UniProt: the universal protein knowledgebase in 2021. Nucleic Acids Res, 49(D1), D480–D489. https://doi.org/10.1093/nar/gkaa1100

58. Vogt, N. M., Kerby, R. L., Dill-McFarland, K. A., Harding, S. J., Merluzzi, A. P., Johnson, S. C., Carlsson, C. M., Asthana, S., Zetterberg, H., Blennow, K., Bendlin, B. B., & Rey, F. E. (2017). Gut microbiome alterations in Alzheimer’s disease. Sci Rep, 7(1), 13537. https://doi.org/10.1038/s41598-017-13601-y

59. Vrba, J., Kren, V., Vacek, J., Papouskova, B., & Ulrichova, J. (2012). Quercetin, quercetin glycosides and taxifolin differ in their ability to induce AhR activation and CYP1A1 expression in HepG2 cells. Phytother Res, 26(11), 1746–1752. https://doi.org/10.1002/ptr.4637

60. Wang, M., Li, Y. J., Ding, Y., Zhang, H. N., Sun, T., Zhang, K., Yang, L., Guo, Y. Y., Liu, S. B., Zhao, M. G., & Wu, Y. M. (2016). Silibinin Prevents Autophagic Cell Death upon Oxidative Stress in Cortical Neurons and Cerebral Ischemia-Reperfusion Injury. Mol Neurobiol, 53(2), 932–943. https://doi.org/10.1007/s12035-014-9062-5

61. Wei, X. J., Wu, J., Ni, Y. D., Lu, L. Z., & Zhao, R. Q. (2011). Antioxidant effect of a phytoestrogen equol on cultured muscle cells of embryonic broilers. In Vitro Cell Dev Biol Anim, 47(10), 735–741. https://doi.org/10.1007/s11626-011-9464-x

62. Wiatrak, B., Kubis-Kubiak, A., Piwowar, A., & Barg, E. (2020). PC12 Cell Line: Cell Types, Coating of Culture Vessels, Differentiation and Other Culture Conditions. Cells, 9(4). https://doi.org/10.3390/cells9040958

63. Wicker, J., Lorsbach, T., Gutlein, M., Schmid, E., Latino, D., Kramer, S., & Fenner, K. (2016). enviPath--The environmental contaminant biotransformation pathway resource. Nucleic Acids Res, 44(D1), D502–508. https://doi.org/10.1093/nar/gkv1229

64. Wishart, D. S., Tian, S., Allen, D., Oler, E., Peters, H., Lui, V. W., Gautam, V., Djoumbou-Feunang, Y., Greiner, R., & Metz, T. O. (2022). BioTransformer 3.0-a web server for accurately predicting metabolic transformation products. Nucleic Acids Res, 50(W1), W115–W123. https://doi.org/10.1093/nar/gkac313

65. Yang, G., Hong, S., Yang, P., Sun, Y., Wang, Y., Zhang, P., Jiang, W., & Gu, Y. (2021). Discovery of an ene-reductase for initiating flavone and flavonol catabolism in gut bacteria. Nat Commun, 12(1), 790. https://doi.org/10.1038/s41467-021-20974-2

66. Yousofshahi, M., Lee, K., & Hassoun, S. (2011). Probabilistic pathway construction. Metab Eng, 13(4), 435–444. https://doi.org/10.1016/j.ymben.2011.01.006

67. Yousofshahi, M., Manteiga, S., Wu, C., Lee, K., & Hassoun, S. (2015). PROXIMAL: a method for Prediction of Xenobiotic Metabolism. BMC Syst Biol, 9, 94. https://doi.org/10.1186/s12918-015-0241-4

68. Zhang, H., & Tsao, R. (2016). Dietary polyphenols, oxidative stress and antioxidant and anti-inflammatory effects. Current Opinion in Food Science, 8, 33–42.

69. Zheng, S., Zeng, T., Li, C., Chen, B., Coley, C. W., Yang, Y., & Wu, R. (2022). Deep learning driven biosynthetic pathways navigation for natural products with BioNavi-NP. Nat Commun, 13(1), 3342. https://doi.org/10.1038/s41467-022-30970-9

