## Supplementary figures and images for "A Chemical Reaction Similarity-Based Prediction Algorithm Identifies the Multiple Taxa Required to Catalyze an Entire Metabolic Pathway of Dietary Flavonoids"

### Supplementary Figure S1. Example input and output

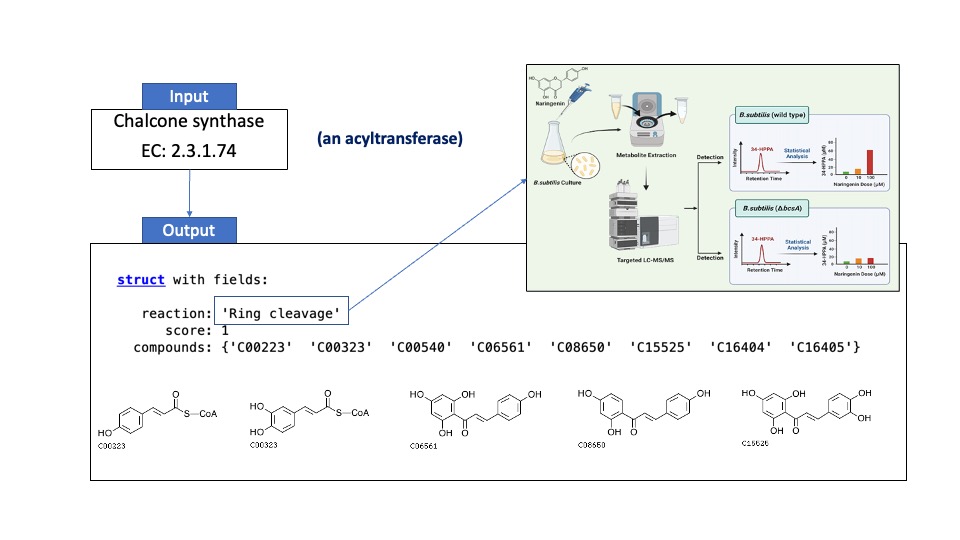

### Supplementary Figure S2. Concentrations of possible microbial metabolites of apigenin and naringenin detected in the apigenin treated B. animalis mono

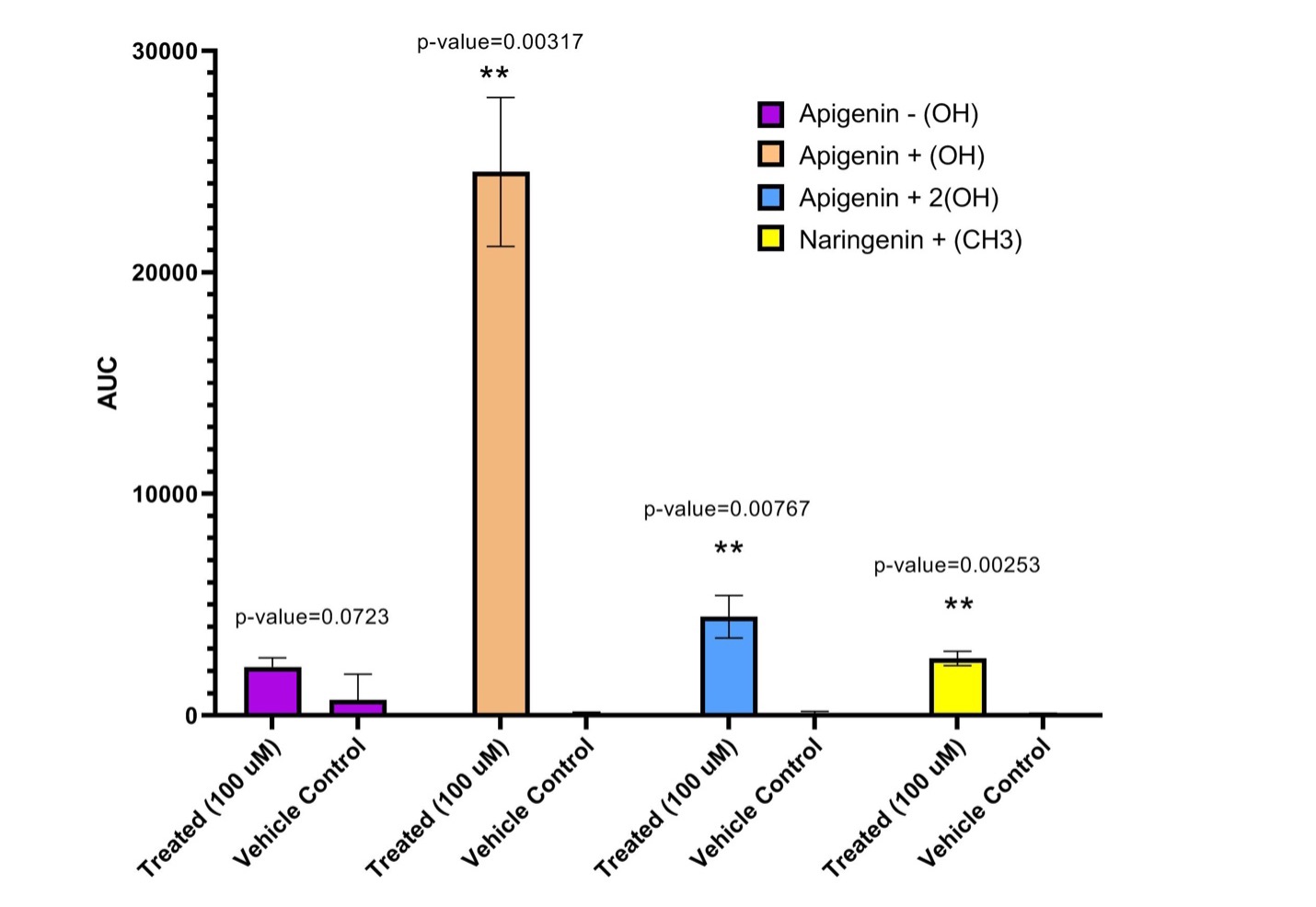
